# Surface expressed *Plasmodium* circumsporozoite protein (CSP) modulates cellular flexibility and motility

**DOI:** 10.1101/2021.08.04.455043

**Authors:** Aditya Prasad Patra, Vrushali Pathak, Segireddy Rameswara Reddy, Aditya Chhatre, Crismita Dmello, Satya Narayan, Dipti Singh, Kota Arun Kumar, Sri Rama Koti Ainavarapu, Shobhona Sharma

**Author notes:** To whom correspondence should be addressed. (S.S.), (S.R.K.A.).

## Abstract

*Plasmodium falciparum* circumsporozoite protein (CSP) is a critically required abundant surface protein of sporozoites and a major vaccine candidate. However, neither the structure nor the role of CSP in sporozoite motility is well understood. Our recent *in vitro* data, from single-molecule pulling experiments suggested a mechanically pliable structure for *P. falciparum* CSP. By engineering vegetative cells of the cellular slime-mold *Dictyostelium discoideum* with regulatable CSP surface expression, we report evidence for direct involvement of CSP towards conferring elastic properties and motility of the cells. With an increase in the surface-CSP levels by 5–8-fold, the Youngs moduli of the cells, observed through atomic force microscopy, decreased around 2-fold, with a concomitant increase in motility by about 2-fold. Interestingly, only full length CSP expression conferred maximal flexibility and motility, as opposed to repeat region alone or the flanking domains of CSP. The enhanced motility of the CSP-expressing cells was abrogated with anti-CSP antibodies as well as phospholipase cleavage of CSP, indicating specific contribution of CSP towards motility. Measurements of the Youngs moduli of *Plasmodium berghei* midgut (MG) and salivary gland (SG) sporozoites revealed an inverse correlation with CSP levels with a decrease from 1.1 kPa to 0.3 kPa as the CSP concentration doubled from MG to SG sporozoites. We hypothesize that high CSP level lowers the stiffness of sporozoites possibly through its pliable surface-coat, leading to cellular flexibility. These findings may explain a sporozoites developmental ability to enhance its CSP levels during transition from midgut to salivary glands to suit a migratory mode in the host, needed for successful hepatocyte invasion.

## Introduction

Malaria remains a major scourge throughout the tropical world. Currently the most effective trial vaccine against malaria, RTS,S, includes domains from only one *Plasmodium falciparum* malaria antigen- the circumsporozoite protein (CSP) (1). CSP is the major surface protein on sporozoites, and in all species of *Plasmodia*, they exhibit the same structural organization with a unique species specific central repeat region (2,3). Certain reversible conformational changes of CSP are required for infection of the mammalian host (4), but detailed structural and dynamic properties of CSP remain incompletely understood. Earlier studies have postulated a superhelical structure for the repeat regions (5), but using single-molecule force spectroscopy (SMFS), we have recently demonstrated that the repeat regions of CSP are heterogeneous and largely unstructured (6). A well-folded protein domain requires about 200 pico-Newton (pN) to unwind such domains, however nearly 40% of the CSP-repeat molecules required <10 pN to unfold, while the remaining 60% of the population consisted of partially folded structures with low mechanical resistance (70 pN). SMFS of CSP molecules exhibited two major conformational populations (6), possibly indicating the open and collapsed forms, consistent with earlier observations (4).

CSPs from different *Plasmodium* species do not exhibit sequence homology except in two small regions viz., region I, which is immediately preceding the tandem repeat region, and the other in the C-terminal region, which is homologous to the thrombospondin type-1 repeat (TSR) superfamily (2,3). The extreme C-terminus contains a canonical glycosyl-phosphatidyl inositol (GPI) anchor addition sequence, although the presence of a GPI anchor has not been demonstrated directly. Both complete *CSP* KO, and mutants exhibiting diminished levels of CSP, revealed a critical role of the protein in sporulation within oocyst (7,8). In an attempt to further appreciate the structure-function aspects of CSP, several deletional mutants were generated that lacked the N-terminus, the central repeat region and the canonical GPI anchor sequence. With an exception to N-terminal domain deletion, other mutants resulted in defective sporoblasts (9–11). The N-terminal deletion mutant yielded sporozoites, but they failed to migrate and infect hepatocytes effectively (11,12). The diverse repeat regions of all CSPs are located centrally and are immunodominant (13) and could be involved in immune evasion (14–16). Regions I and the C-terminal TSR of CSP are postulated to be involved in salivary gland and liver heparin sulfate proteoglycan receptor recognition sites, respectively (11,17–19).

Sporozoites undergo an unusually long journey following their egress from oocyst. Within mosquito hemocoel, they migrate to the salivary glands and wait for transmission. Once introduced into skin of the mammalian host (20), they traverse through the cells of the dermis, tissue matrices, and vascular endothelial cell junctions prior to their selective arrest in liver cells. The random entry and exit of sporozoites through cells, referred to as migration, is a non-passive process that activates the sporozoites in the presence of albumin and is critically time dependent (21). Within liver, Kupffer cells have been shown to be the portal of entry by forming transient vacuoles (22,23). Following exit from Kupffer cells, the sporozoites move through couple of hepatocytes prior to switch into productive mode of invasion, accompanied by formation of an intrahepatic parasitophorous vacuole (24,25). Intravital microscopy has revealed that the sporozoites move fast in the blood flow (1-3 μm/s) and can be seen to squeeze through Kupffer cells, endothelial cells and hepatocytes (21,26), and clearly spend great energy towards effective migration, exhibiting various forms of movements such as circular, linear and meandering (12,27,28). The motility of sporozoites is fueled by an acto-myosin molecular motor present in the cortical region of the sporozoite, underneath the plasma membrane. Multiple components like gliding associated proteins, adaptor proteins, and the thrombospondin-related anonymous protein (TRAP) (29–31) act in concert to facilitate a substrate dependent motility referred to as gliding motility. Despite its abundance, there is no evidence that CSP is a constituent of the molecular motor complex, and hence its role in cell motility remains unclear. Considering the high speed of sporozoite migration in vivo, it is likely plausible that the sporozoites may experience considerable frictional and drag forces that may impose constraints on its motility. Based on previous observation by SMFS (6), we hypothesize that the densely coated CSPs provide a structurally pliable cushion cover on the sporozoite surface that likely lowers the stiffness of the cells, provides lubrication and eases cell motility and traversal.

To test this hypothesis, an approach to manipulate CSP levels on the sporozoite surface would be ideal. However, it is not practically feasible, as complete knock-out of *CSP* gene negatively impacts sporozoite development while abrogation of normal levels of CSP affects their structural integrity, yielding hypomorphs (7,8). Therefore, in order to assess the contribution of varying levels of surface expressed CSP towards stiffness and motility *in vivo*, we used an amoebic stage of the cellular slime mold *D. discoideum* as a model system. Anchoring and characterization of *P. falciparum* CSP on the surface of *D. discoideum* cells has been reported earlier (32). In the current study, using a combination of an atomic force microscope (AFM) and video microscopy, we successfully measured respectively the stiffness (Young’s modulus) and motility of the amoebic cells expressing on its surface, either the full length CSP or its deletion variants. Further, we compared the stiffness of the CSP-expressing *D. discoideum* cells with *P. berghei* sporozoites and observed the values to be in the same order, at about 1 kPa. We also observed a direct correlation between surface levels of the full-length CSP and a reduction in stiffness, along with an enhancement in the motility of the cells.

## Results

### Expression of CSP on the *D. discoideum* cells

The two CSP plasmid-constructs used in this study were Periv CS and Pac15 CSP (kind gifts from Prof. N. Fasel) (32). The CSP expression in Periv CSP and Pac15 CSP was driven respectively by Ras and actin-15 promotor (Fig. S1). In each of the constructs, the N-terminal 18 aa domain (CSP leader) and the C-terminal 23 aa (GPI-anchor) of *PfCSP* were replaced respectively with a 26 aa leader sequence and a 48 aa stretch of GPI anchor from *Dictyostelium* Contact Site A antigen (CSA) (Fig. S1). Transfected *Dictyostelium* cells expressed CSP soluble/cellular and secreted forms of protein (Fig. S2), as demonstrated earlier (32).

### Stiffness measurement of *D. discoideum* cells expressing surface-CSP

The stiffness (Young’s modulus) of a living cell is usually calculated from the force–displacement curves measured by indentation tests. Most common analysis models for cell indentation are based on the Hertz theory, and the stiffness of *D. discoideum* cells expressing surface-CSP was measured with the help of AFM-indentation assays (33). Schematic representations of the AFM used for the cell-indentation assays, the area of the amoebic cell probed by the AFM cantilever, and the force vs distance traces obtained upon such probing are shown in Fig.1A-C. A representative trace result of cell indentation assay showing force vs distance graph for AX2 cells is presented in Fig.S3. Representative force vs distance traces for the transfected cells are shown in Fig. S3. The control *D. discoideum* AX2 cells showed considerable variation in the Young’s modulus. The average Young’s modulus was observed to be about 12 kPa which was comparable to the value reported earlier (33). The variation could be in the values within each cell, or between different cells. An analysis of all points of observation and the mean value per cell were plotted, but no significant differences were observed with the two ways of analysis (Fig. S4), indicating that there were no significant differences in Young’s modulus measured for each cell, however, in the vegetative phase, the cells were heterogenous in their stiffness. Therefore, average stiffness values calculated for individual cells have been used for all subsequent *D. discoideum* dot plots. As reported earlier, *D. discoideum* cells were starved before adding appropriate concentration of cAMP to induce the promoter activity and hence the expression of CSP (32–34). We observed that starvation itself caused the Young’s modulus of AX2 cells to decrease significantly from 7.3 to 2.7 kPa, but the CSP-transfected cells exhibited a further decrease in Young’s modulus over and above the decrease in the control AX2 cells (Fig. S5). Since starvation induces internal cAMP production in *D. discoideum* cells, which may have a confounding effect on the CSP produced, subsequent assays were carried out in the presence of cAMP without starvation. The induction of transfected CSP-expressing cells with 400 μM cAMP resulted in a 3-6 fold reduction in Young’s modulus as compared to the AX2 cells treated with 400 μM cAMP (Fig. 1E).

**Fig. 1.**
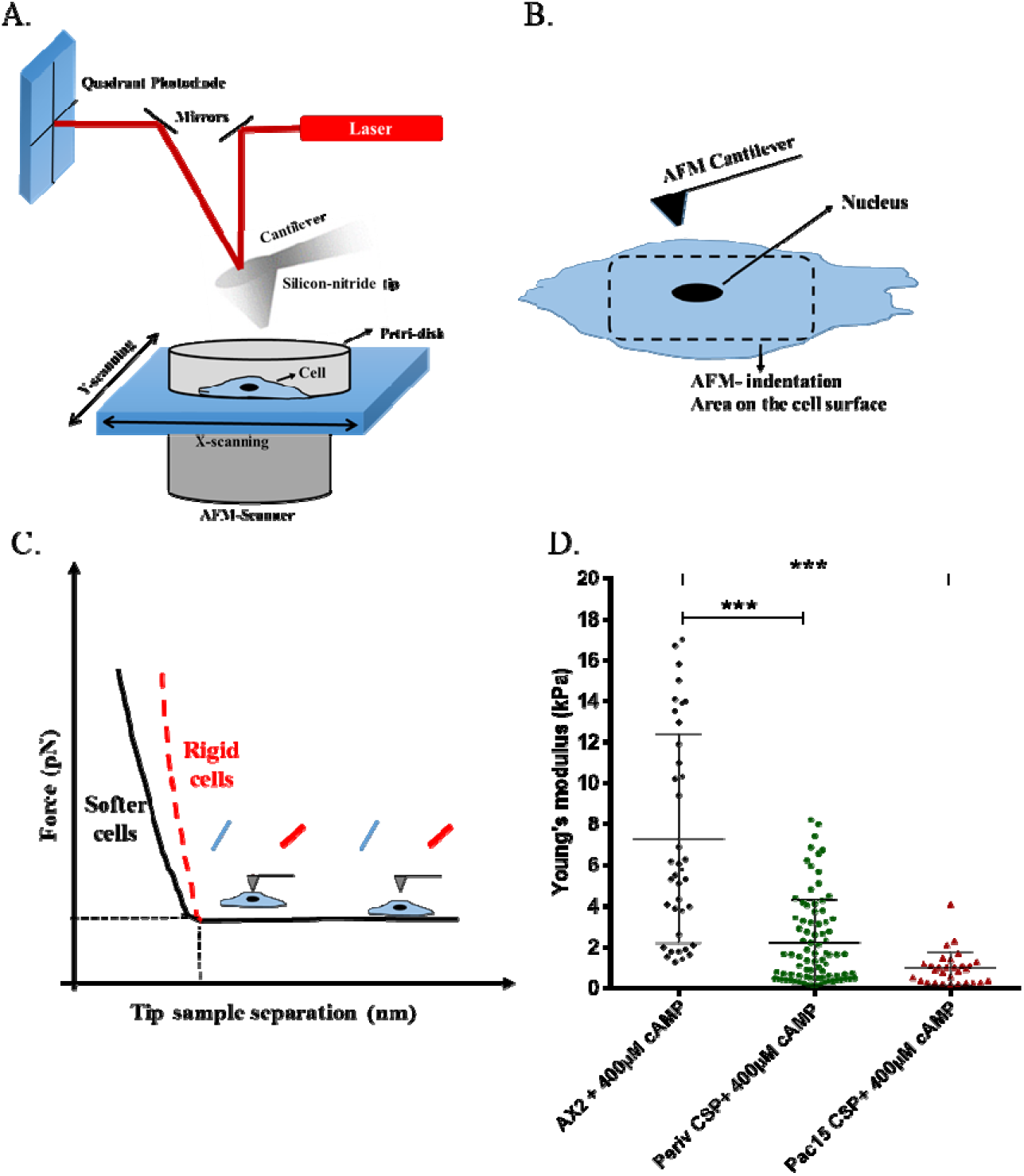
Cell-indentation assays. A) Schematic representation of the AFM set up used for the cell-indentation assays to measure the stiffness of the cells. Polylysine coated petri-dish was used for adhering cells as described in methods section. B) Cartoon diagram showing the indentation area on a cell. C) Cartoon diagram of force vs distance traces for stiffer and softer cells. D) Stiffness measurements of *D. discoideum* cells under different conditions: AX2 (control) cells with and without cAMP; Periv-CSP and Pac15-CSP represent AX2 cells transfected with Periv-CSP and Pac15-CSP constructs. \**p*<0.05; \*\*\**p*<0.001.

In order to establish surface expression of CSP on the transfected cells, flow cytometric analysis was performed using anti-(NANP)_5_ antibodies. Representative plots of mean fluorescent intensity (MFI) are shown in Fig. 2A, whereas a compilation of data from multiple experiments is shown in Table S1. A significant amount of surface-CSP was detected even without the addition of cAMP, indicating a basal rate of CSP synthesis in the transfected cells. As the cAMP concentration was increased from 50 to 400 μM, the MFI increased about 5-fold for Periv-CSP, and about 2-fold for Pac15-CSP (Fig. 2A, Table S1). The flow cytometry results revealed the presence of CSP on the cell surface and the feasibility of altering the levels of surface expression of CSP by varying the cAMP concentration. The control AX2 cells did not show any labeling with anti-(NANP)_5_ antibodies under these conditions (data not shown). The Young’s moduli of the cells showed an inverse correlation with the surface-CSP levels (Fig. 2B-C, Table S1). With an increase in the surface-CSP by about 5-8 fold, there was a 3-fold decrease in the Young’s moduli of the amoebic cells (Fig. 2B-C, Table S1). The results demonstrated a direct correlation between levels of CSP expression on the amoebic cell surface with a decreased stiffness of the cells.

**Fig. 2.**
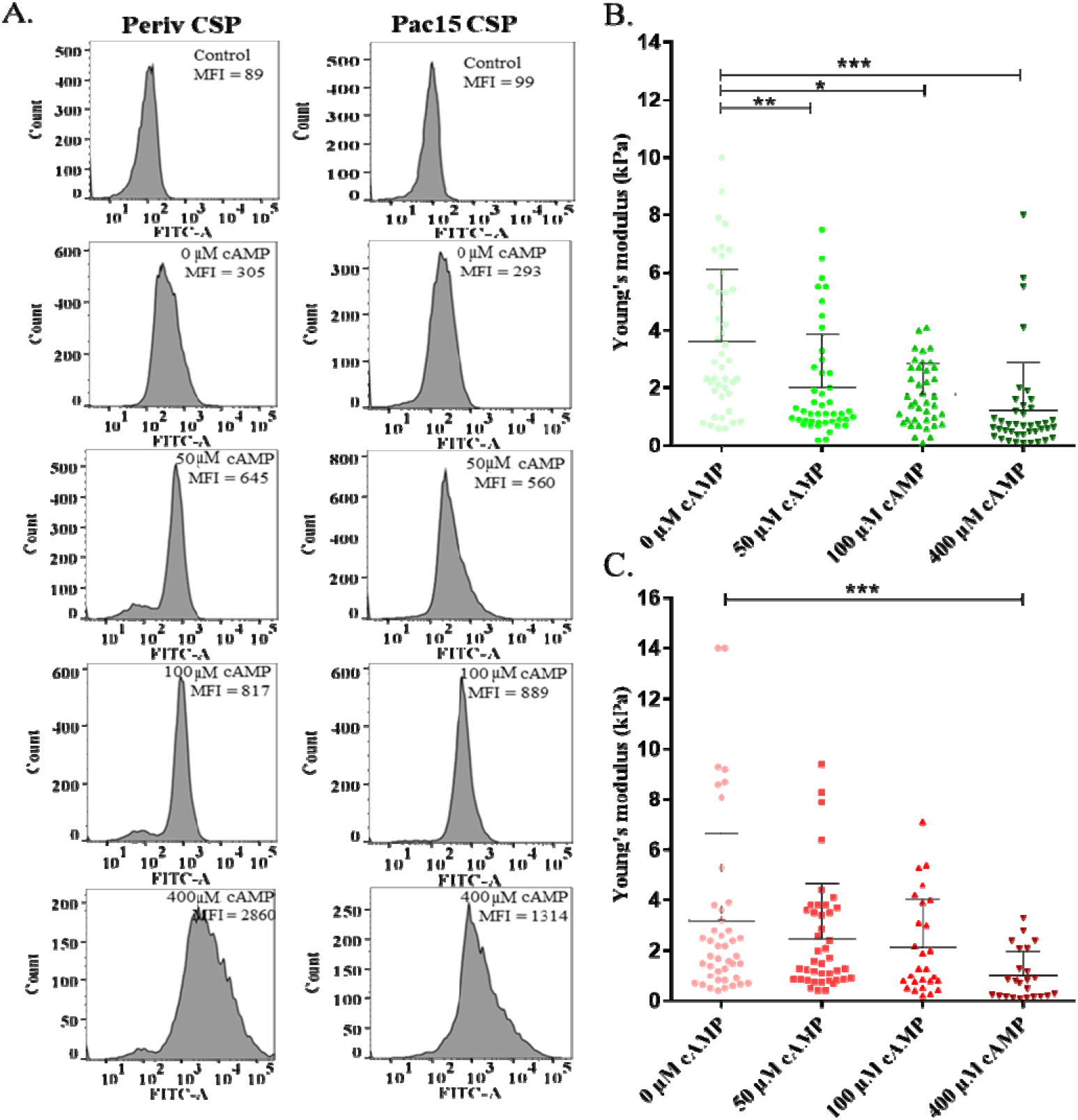
CSP concentration dependence of cell stiffness. Surface expression of CSP determined by flow cytometry and stiffness measured by AFM indentation assay. A) Representative images of mean fluorescence intensity (MFI) of surface-CSP expression on Periv-CSP and Pac15-CSP transfected *D. discoideum* cells induced by incubation with different concentrations of cAMP. The vertical panels show secondary antibody control, 0 μM cAMP, 50 μM cAMP, 100 μM cAMP and 400 μM cAMP treated cells, stained with anti-(NANP)_5_ antibodies. B) and C) Concomitant stiffness of the *D. discoideum* cells transfected with Periv-CSP and Pac15-CSP plasmids, respectively, and induced with cAMP. \**p*<0.05; \*\**p*<0.01; \*\*\**p*<0.001.

### Motility of *D. discoideum* cells expressing surface-CSP

To measure the motility of the *D. discoideum* cells expressing CSP, the movement of the cells through agarose matrix were video imaged, using a chemotactic set up with folate as the chemo-attractant (Fig. 3A). A representative video track is shown in Fig S6. The speed of the control cells AX2 was observed to be 8.1 ± 1.5 μm/min (Fig. 3B), similar to that reported earlier (35). However, upon treatment with 400 μM cAMP, *D. discoideum* cells showed a significant enhancement of speed to those of 18.5 ± 2.8 μm/min and 19.5 ± 3.8 μm/min for Periv-CSP and Pac15-CSP cells, respectively (Fig. 3B). When the transfected *D. discoideum* cells were subjected to increasing concentrations of cAMP (50, 100, and 400 μM cAMP), the motility showed a graded increase by 25 to 65% (Fig. 3C-D, Table S1).

**Fig. 3.**
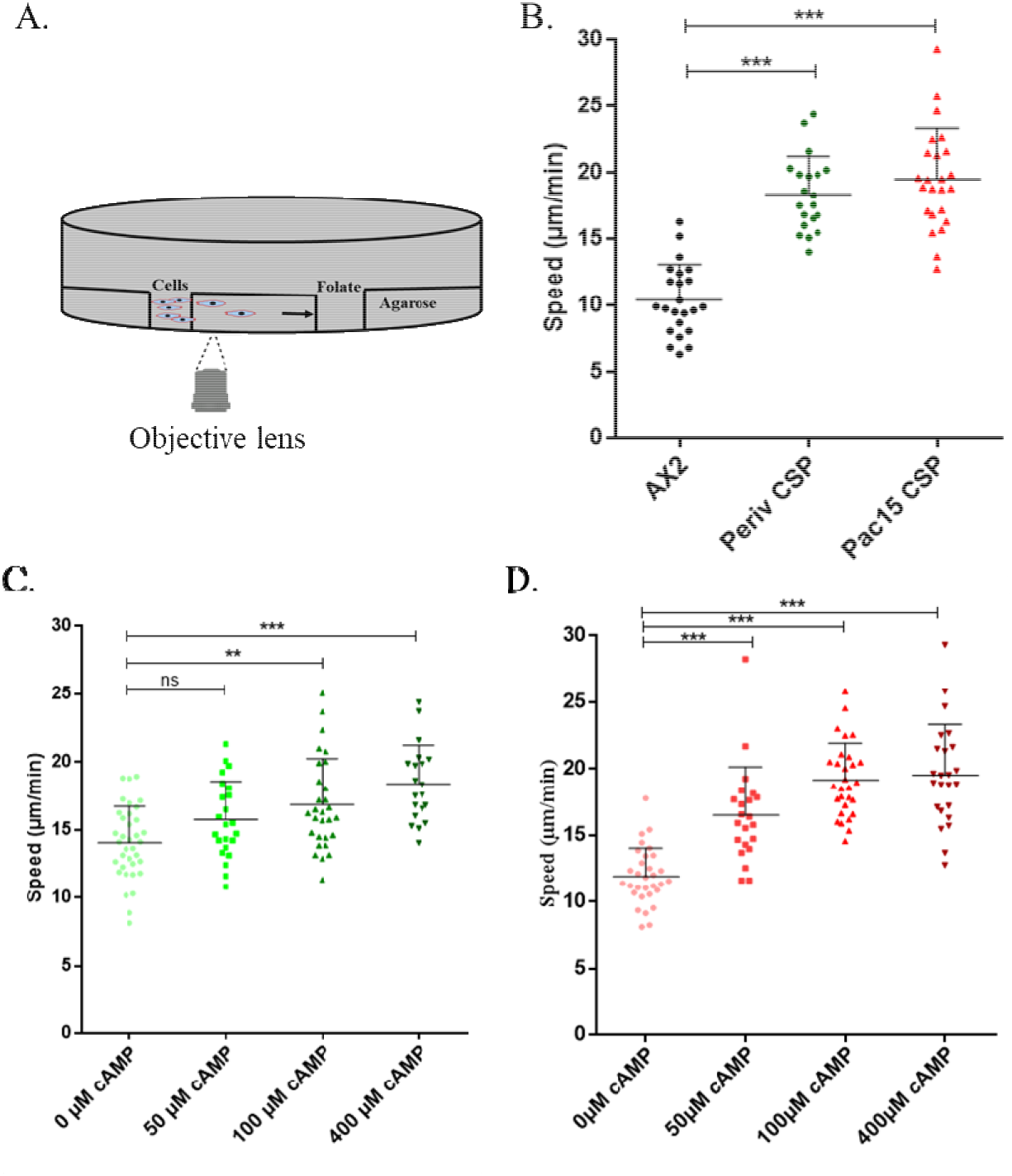
Cell motility of transfected *D. discoideum* cells. A) Schematic representation of the agarose plate with folate as the chemoattractant. The speed of the moving cells was determined using time-lapse video. B) Dot plots showing the speed of AX2 (control) cells, and AX2 cells transfected with Periv-CSP and Pac15-CSP constructs induced with 400 μM cAMP. C) and D) Dot plots of speed of *D. discoideum* cells transfected with C) Periv-CSP and D) Pac15-CSP induced with different concentrations of cAMP. \**p*<0.05; *\*\*p*<0.01; *\*\*\*p*<0.001.

To test whether the enhanced motility was specific for CS-protein, CSP-expressing *D. discoideum* cells were incubated with anti-(NANP)_5_ antibodies (Fig. 4). Significant inhibition in the motility of the cells were observed, both with increase in the time of incubation and the antibody titer (Fig. 4), indicating a dependence of CSP in cell motility. However, the immobilization could be due to cross linking of CSP and the anti-CSP antibodies. A better way of checking specificity would be to remove CSP and test for cell motility. Also, since the CS-proteins were tagged with *D. discoideum* CSA-GPI anchor, we envisaged that with an enhancement in the surface-CSP levels, there would be a concomitant increase in the CSA-GPI anchor moieties. Since GPI moieties contribute to enhanced stiffness through raft micro-domains (36)(36), our observations could not rule out the possibility of enhanced motility being attributable mainly to increased levels of CSA-GPI anchor. In order to test a dominating role of CSA-GPI anchor in mediating cell motility, assays were set up that involved cleavage of CSP from the CSA-GPI anchor using GPI-Phospolipase-D (GPI-PLD) (37). A time course of GPI-PLD treatment revealed a dramatic depletion of surface-CSP within 5 minutes of treatment, with a concomitant decrease in cell-motility to very low levels (3-7-fold) (p<0.01; Fig. 5A). However, within 15 minutes the CSP started replenishing the surface and there was recovery in the motility rates (Fig. 5A-C). Since cleavage using GPI-PLD only released CSP, without affecting CSA-GPI levels, we conclude that the reduced motility of the cells after GPI-PLD treatment relates specifically to the loss of surface-CSP molecules.

**Fig. 4.**
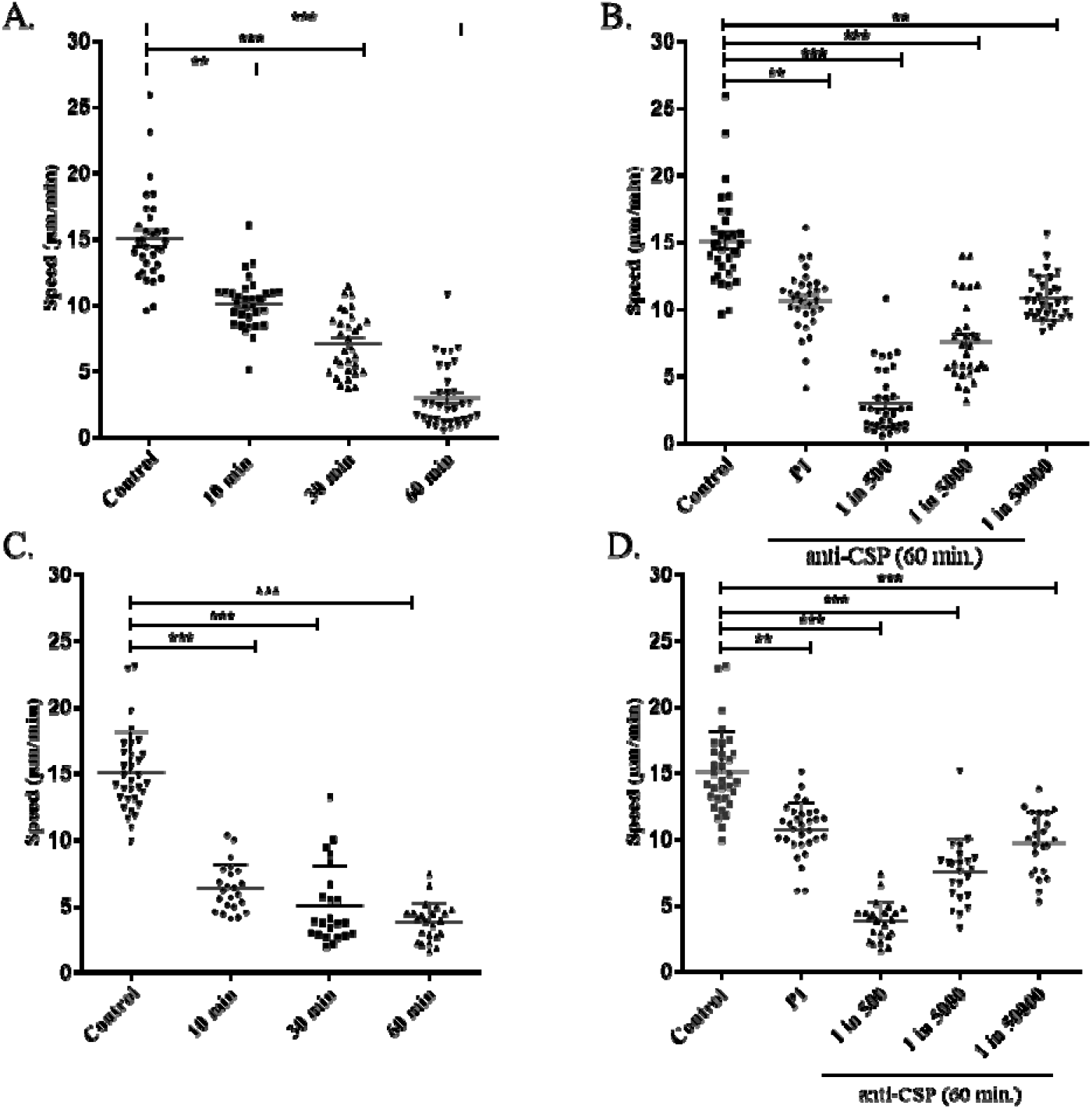
Speed of transfected *D. discoideum* cells in the presence of anti-PfCSP antibody. Dot plots of speed of transfected *D. discoideum* cells induced with 400 μM cAMP. A) and C) *D. discoideum* cells transfected with Periv-CSP and Pac15-CSP constructs, respectively, incubated with anti-PfCSP antibodies for different duration. B) and D) Dot plots of speed of *D. discoideum* cells transfected with Periv-CSP and Pac15-CSP constructs, respectively, treated with different titres of the anti-PfCSP antibodies. *\*\*p*<0.01; *\*\*\*p*<0.001.

**Fig. 5.**
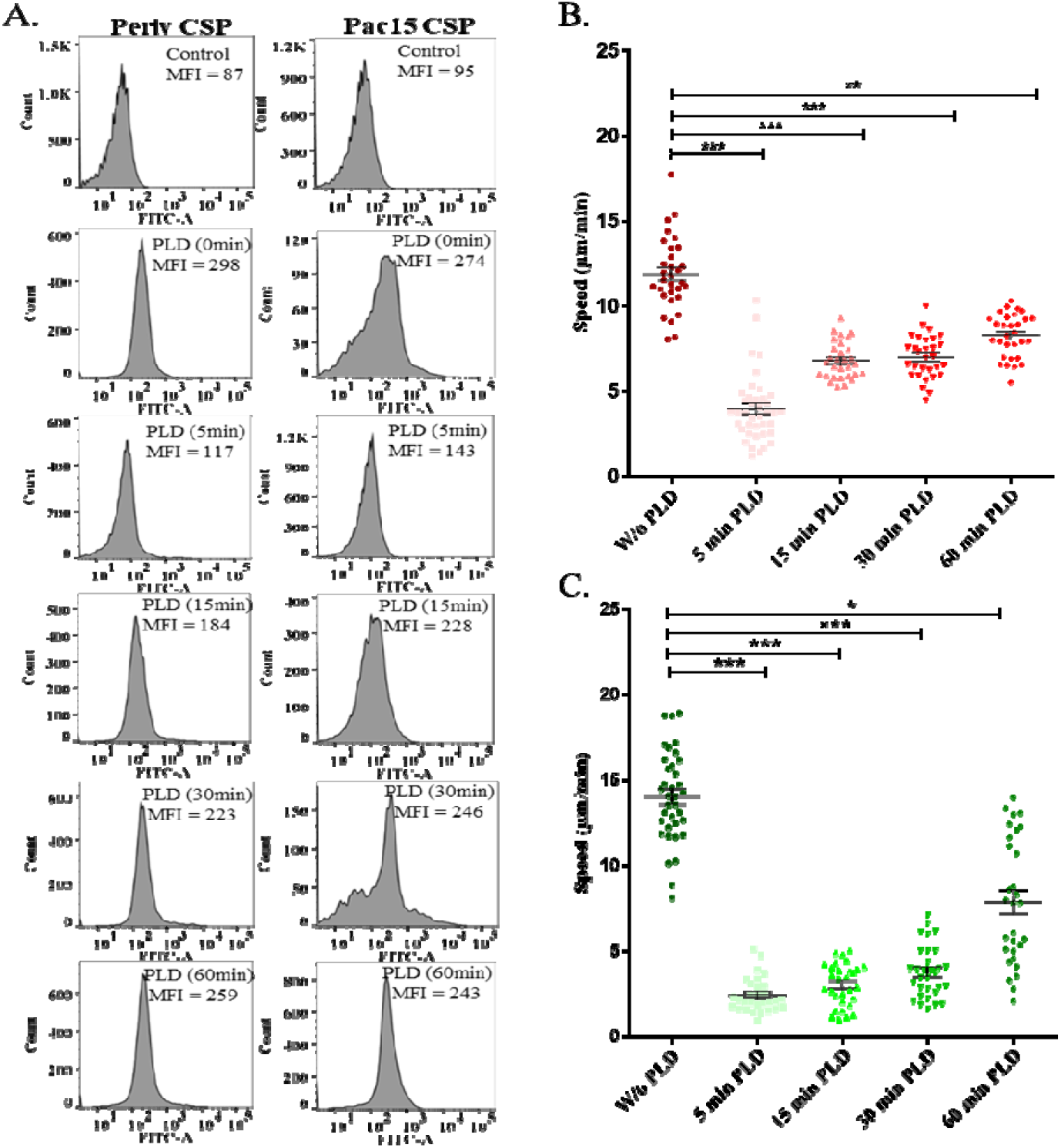
Speed of transfected *D. discoideum* cells treated with Phospolipase D. A) Flow cytometry data showing the MFI of CSP expression on Periv-CSP and Pac15-CSP transfected *D. discoideum* cells at 0 μM cAMP., upon treatment with GPI-Phospolipase D (GPI-PLD) for different duration. Dot plots of speed of the *D. discoideum* cells transfected with B) Periv-CSP and C) Pac15-CSP constructs upon treatment with GPI-PLD for different duration. \**p*<0.05; *\*\*p*<0.01; *\*\*\*p*<0.001.

### Stiffness and motility of *D. discoideum* cells expressing sub-domains of CSP

In order to further explore the functions of different domains of CSP, Periv constructs containing sub-domains of CSP were generated (Fig. S1) and assayed for stiffness and motility (Fig. 6; Figs. S6-S9). As shown in Fig. S1, the CSP repeat domain protein (Periv CSP-R), CSP-protein without the N-terminal domain (Periv CSP-Ndel) and CSP-protein with just the C-terminal domain (Periv CSP-NRdel) were constructed in Periv vector using the same promoter region, signal sequence and GPI anchor. The construct of just the N-terminal domain (CSP-RCdel) was also attempted but we were not successful in obtaining a stable clone. For comparison with a fairly rigid protein of a size similar to CSP, four repeats of the I27 domain from the I-band of the giant muscle protein titin (38) was also cloned in the same vector, resulting in Periv (I27)4. The protein expressions could be induced using cAMP as measured through flow cytometry analysis (Fig. S7). Some amount of leaky expression was observed in all the cases, but the expression levels were comparable to full-length Periv-CSP cells at the 400 μM cAMP induction. The Young’s moduli values in the absence and presence of cAMP for these cells are shown in Fig. S8. The cAMP induced cells were subjected to stiffness measurements and motility assays (Fig. 6 B, C). The Young’s moduli for the CSP-R and CSP-Ndel expressing cells were not significantly different from the full-length CSP (Fig. 6 B). Each of these constructs contained the repeat-region, and therefore the flexibility could be provided by the repeat domain of CSP. The C-terminal region (CSP-NRdel), containing the structured thrombospondin like domain (39) and the well-folded (I27)4 (38) expressing cells were significantly more rigid (Fig. 6 B).

**Fig. 6.**
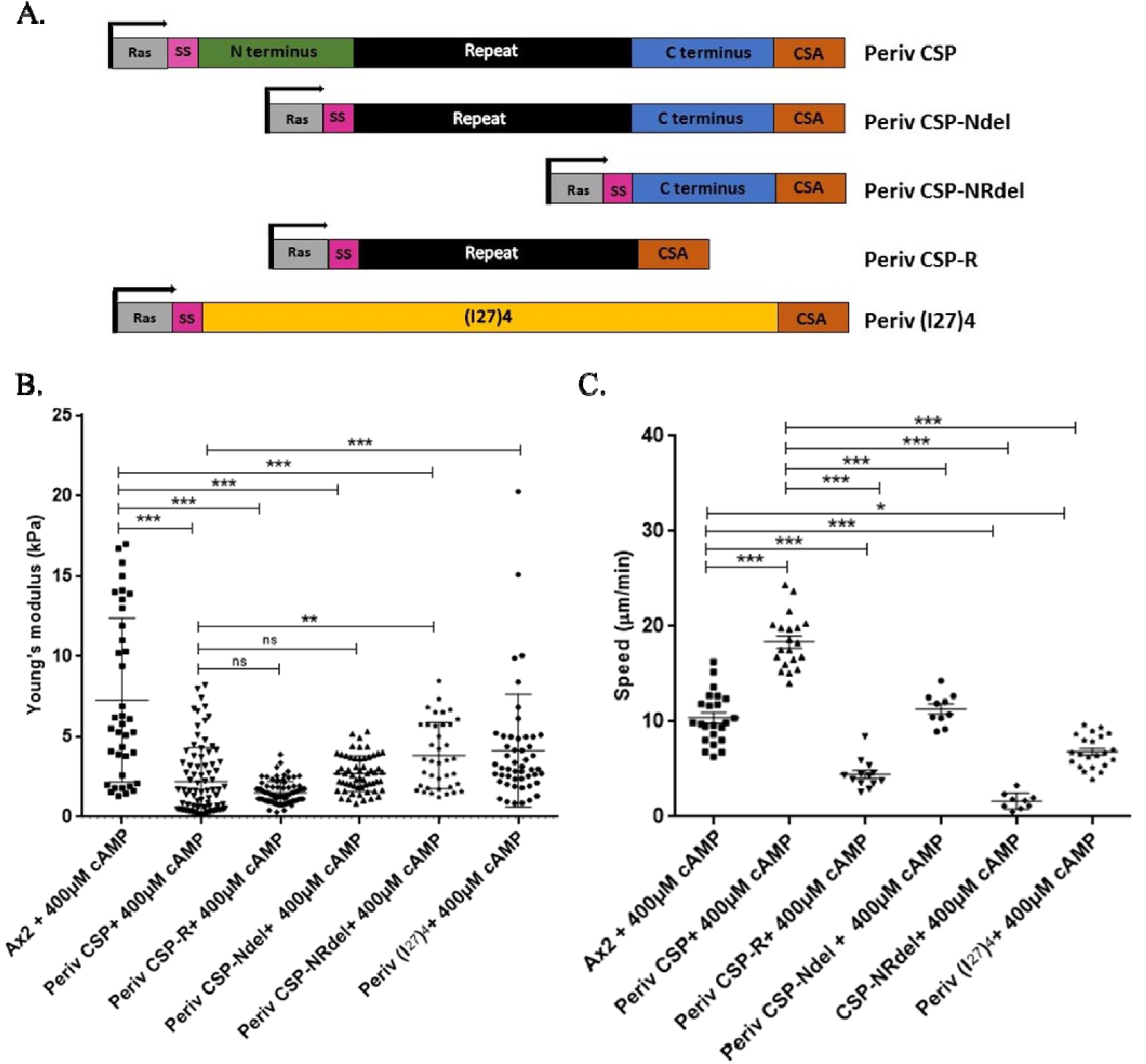
Cell stiffness and motility of *D. discoideum* expressing sub-domain of CSP and (I27): A) Cartoon representation of different subdomain of CSP and (I27)_4_ were cloned into Periv vector under Ras promoter. The expression of the surface expressed CSP subdomain control by cAMP induction. Dot plots of B) cell stiffness and C) speed of ****D. discoideum**** cells expressing different subdomain of CSP and (I27)_4_ induced with 400 μM cAMP. \**p*<0.05; *\*\*p*<0.01; *\*\*\*p*<0.001.

While we hypothesized an inverse correlation between the stiffness of the cells and their motility, surprisingly, the motility of the cells expressing different domains of CSP did not correlate directly with the stiffness. Cells expressing full-length CSP exhibited maximum motility, as opposed to any other sub-domains of the protein (Fig. 6C, Fig. S8, 9). Cells expressing repeat region alone, and CSP with N-terminal deleted showed significantly lower motility as compared to CSP-expressing cells, although these cells exhibited comparable values of Young’s moduli. The more rigid cells, Periv CSP-NRdel and Periv (I27)4, also showed low motility. Each of the CSP-subdomain expressing cells, except CSP-Ndel expressing cells, were less motile than the control AX2 cells, although AX2 cells were stiffer than each of these cell types. We conclude that motility was dependent on parameters other than just the stiffness of the cells. Each of these transfected cells had comparable amounts of surface protein expression but exhibited significantly different motility, supporting our earlier conclusion that the motility of the cells could not be attributed to the CSA-GPI moiety.

### Stiffness of *P. berghei* sporozoites

Expression of full-length CSP on *D. discoideum* cells decreased the cell stiffness to about 1-3 kPa. However, it was important to assess whether the stiffness values of these *D. discoideum* cells were in the physiological range to that of *Plasmodium* sporozoites that are accentuated with a thick coat of CSP. Towards this end, we isolated midgut sporozoites (MG-SPZ) and salivary gland sporozoites (SG-SPZ) of the rodent malaria parasites *P. berghei* and subjected them to measurements of CSP-level and stiffness (Fig. 7). Protein lysates generated from similar number of MG-SPZ and SG-SPZ were resolved on SDS-PAGE, and subjected to immunoblotting. Considerably higher levels of CSP was observed in the SG sporozoites (Fig. 7A). Both immunofluorescence and flow cytometry data demonstrated a 2-fold higher level of surface-CSP in the SG-SPZ as compared to MG-SPZ (Fig. 7B-D). AFM indentation measurements demonstrated about 3-fold decrease in Young’s moduli, from 0.87 ± 0.83 kPa in MG-SPZ to 0.32 ± 0.2 kPa in SG-SPZ, pointing to an inverse correlation between surface-CSP levels and the stiffness (Fig. 7E(i)). The midgut sporozoites exhibited large variations in the Young’s modulus, with some apparent outlier sporozoites, likely attributable to the patchy CSP coating on the MG-SPZ (Fig. 7B) and the choice of the region indented by the AFM cantilever. Apparently, this may have been the case, because when we plotted all the data points, the Young’s moduli values were well dispersed for MG-SPZ (Fig. 7E(ii)). The SG-SPZ Young’s moduli values were better clustered, and the averages with all data points were 0.33 ± 0.23 and 1.13 ± 0.94 for the SG-SPZ and MG-SPZ, respectively. It is to be noted that the Young’s moduli of *D. discoideum* cells expressing surface-CSP at 400 μM cAMP (1-2 kPA) were comparable to that of *P. berghei* sporozoites (0.3-1.1 kPa), validating the choice of transfected *D. discoideum* cells as a surrogate model for comparison of stiffness and motility measurements of sporozoites.

**Fig. 7.**
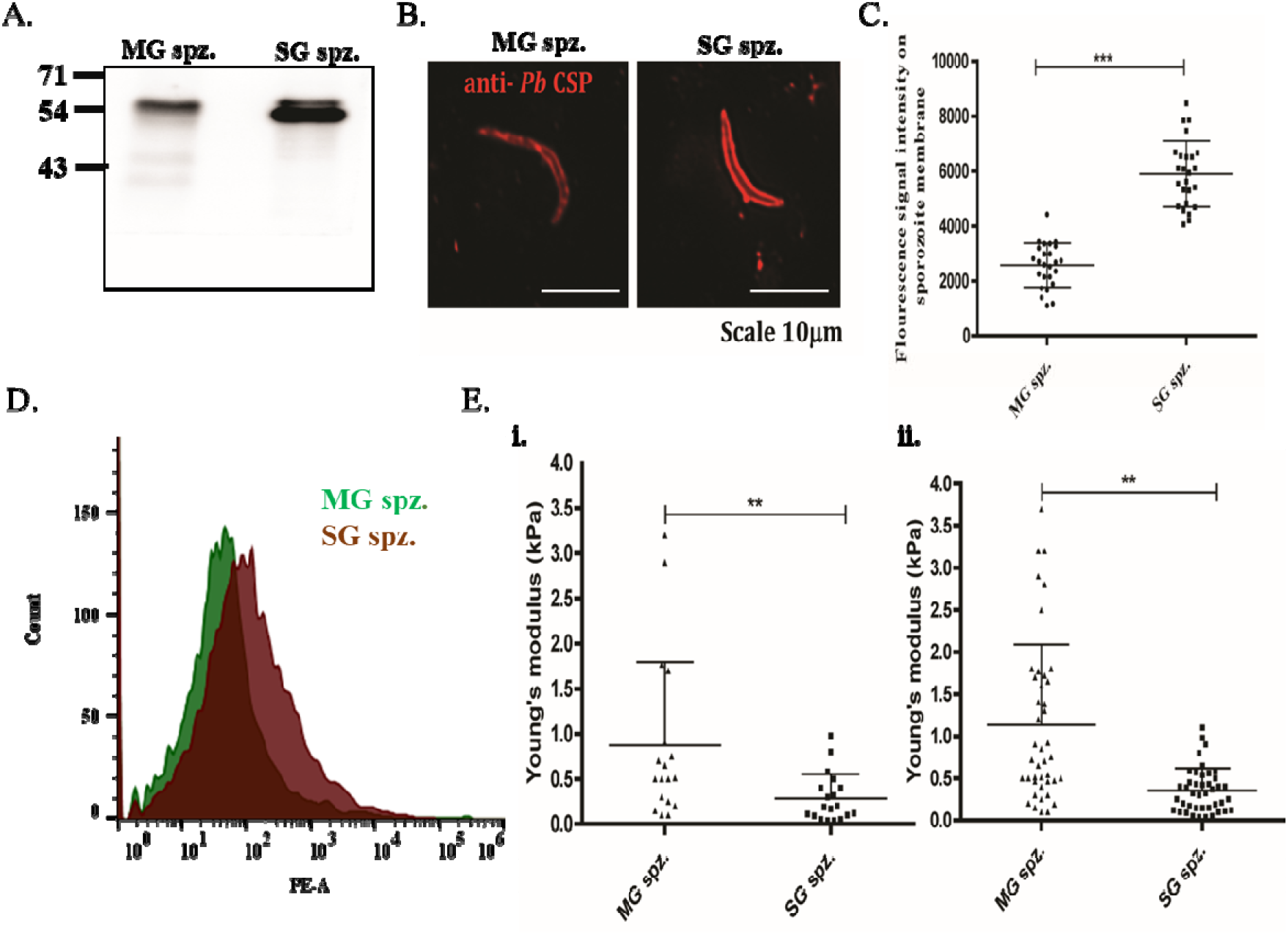
CSP content and stiffness of midgut (MG) and salivary gland (SG) sporozoites of *P. berghei.* A) Western blot of total lysates from 30,000 sporozoites each from midgut and salivary glands; B) and C) Immunofluorescence assay (IFA) of sporozoites probed with anti-PbCSP antibodies. B) Representative image and C) Dot plot showing fluorescence intensity from IFA images of sporozoites. D) Flow cytometry data of sporozoites measured using anti-PbCSP antibodies. E) Dot plot of measured stiffness of MG and SG sporozoites. All individual data points (i) and their mean values per cell (ii) are shown. *\*\*p*<0.01; *\*\*\*p*<0.001.

## Discussion

It has been documented that there is a correlation between cell stiffness and motility (40). Several cancer cells, in particular those exhibiting metastasis, become several-fold softer when measured by various methods, including AFM (40,41). Like metastatic cancer cells, the *Plasmodium* sporozoites also undertake long journeys through tissue matrices and cell junction barriers. However, no information exists regarding the cellular stiffness of *Plasmodium* sporozoites or the role of surface CSP in this stiffness. Since manipulations of CSP levels on sporozoites have been technically difficult and have resulted in defective sporozoites (7–10), we used *D. discoideum* vegetative cells, where the surface expression of CSP could be manipulated. Using AFM and video-imaging we demonstrate that the expression of full-length CSP on the cell-surface of *D. discoideum* amoebic cells results in significantly reduced stiffness and increased motility of the cells. Using *D. discoideum* cells expressing an irrelevant protein such as (I27)4, anti-CSP antibody inhibition studies and through the release of the CSP using GPI-PLD, we showed that the increased motility of the CSP-expressing *D. discoideum* cells could be specifically attributed to CSP. We measured the stiffness (Young’s moduli) of *Plasmodium berghei* sporozoites and the values were 1.13 ± 0.94 kPa and 0.33 kPa ± 0.23 kPa for midgut and salivary gland sporozoites, respectively. To our knowledge, this is the first report of stiffness measurements of *Plasmodium* sporozoites.

### Stiffness of *D. discoideum* cells expressing CSP domains

Several attributes of *D. discoideum* qualified it as a suitable model for investigating the contribution of CSP towards the stiffness and motility of *Plasmodium* sporozoites. *D. discoideum* is a soil amoeba, which is often used as a model for the study of cell motility and chemotaxis (42). Individual vegetative cells undergo chemotactic aggregation in response to signaling mediated by cAMP (43). *D. discoideum* cells have been subjected to stiffness measurements and motility assays earlier. The shape of the cell and several molecular complexes, foremost of which are cytoskeletal and adhesion molecules, play a role in determining the mechanical properties of a cell which is motile across a tissue matrix. It is expected therefore that there will be variations in the stiffness across different parts of the cell and in the case of *D. discoideum* cells it was reported that the apical end of the cell is stiffer than the middle and the rear of the cell, and that actin disruption caused cells to become less stiff (33).

Owing to such constraints, all the measurements in the current study were recorded in the middle region of the *D. discoideum* cell, and the average stiffness values for the AX2 cells were comparable to that reported earlier (33). The measurement of cell stiffness was time consuming, with each cell indentation measurement taking several minutes. These technical shortcomings explained why the GPI-PLD treated *D. discoideum* cells could not be subjected to stiffness measurement, as surface CSP levels started to be restored within 5-10 min (Fig. 5). The stiffness of the sporozoites (SG versus MG) also correlated inversely with the CSP levels and the values of the sporozoite stiffness of <1 kPa were comparable to metastatic cells (40). CSP is known to be secreted at the sporozoite apical end, translocated along the sporozoite surface and released at the sporozoite posterior pole (44). Such directional sloughing may further provide cushioning as the sporozoites move forward.

The contribution of different CSP domains to the stiffness values for the respective *D. discoideum* cells appeared as expected. The CSP, CSP-R and CSP-Ndel expressing *D. discoideum* cells, each containing the repeat region, exhibited comparable stiffness values which were lower than the control cells. Our recent data using SMFS indicates that mainly the repeat region is exposed in about 60% population of CSP molecules, and that very little force is required to unfold this domain (6). Thus, the repeat regions of CSP possess virtually no secondary structure and are likely to be very pliable. Such a property of the repeat region may contribute substantially to the lowering of stiffness observed in CSP, CSP-R and CSP-Ndel expressing *D. discoideum* cells. The CSP-NRdel construct contains mainly the CSP C-terminal region which has sequence homology to the thrombospondin type-1 repeat superfamily (TSR) on the cells. This is the only domain of CSP for which crystal structure exists (39). Within different *Plasmodium* species the structure of CSP-TSR is remarkably conserved and constitutes a well-folded compact structure (39). Similarly, the (I27)4 domain constitutes a well folded muscle protein titin domain (38)(38). Thus, cells expressing CSP-NRdel and (I27)4 would be exposing well-folded structured domains which are likely to contribute towards the higher stiffness of the cells (Fig. 6B).

### Motility of sporozoites and the role of CSP

Does CSP play a role in cell motility? Using *Dictyostelium* cells expressing CSP, we have demonstrated that motility of these cells is enhanced in a concentration dependent manner based on the levels of surface-CSP (Fig. 3). That the enhanced motility is attributable to CSP, and not to enhanced levels of GPI anchor on the membrane was demonstrated by cleaving CSP using GPI-PLD (Fig. 5). As far as sporozoite motility is concerned, the strongest evidence that CSP is involved in motility comes from studies using anti-CSP antibody treatments (44–47). However, the evidence for a direct molecular involvement of CSP in the molecular glideosome complex is missing (48). The zoite motility is powered by the glideosome, which contains the combined activity of an acto-myosin motor located beneath the parasite’s plasma membrane and by protein products of specialized secretory organelles named rhoptries, dense granules, and micronemes (28–31). Extensive studies using fluorescent *P. berghei* sporozoites have demonstrated that acto-myosin based multi-step adhesive forces play a dominant role in the motility and that several parasite proteins such as profilin, coronin, TRAP and the TRAP-like proteins also play roles in sporozoite motility. Sporozoite movements have been classified as linear, circular, meandering, and with sharp turns and it is documented that the chemical milieu as well as radiation affect these movements (27,28,49). While most of the zoite forms of other parasites need to glide to their target host/vector cells over a short distance, the *Plasmodium* sporozoites undergo a long and penetrative journey starting from the mosquito mid-gut to the vertebrate liver cells (20–23,25,50,51). CSP is uniquely present in a huge abundance, with its specific structure of a repeat region flanked by two equally large protein domains, only on the surface of *Plasmodium* sporozoites. The presence of a full-length CSP is critically required for a functioning sporozoite (7–9). Thus, it may be postulated that a correlation exists between the uniquely structured full-length CS-protein and the long and tortuous journey of a sporozoite. Results in this paper, demonstrating enhanced motility of *D. discoideum* cells expressing full-length CSP, support such a postulate. It has been reported earlier that the midgut sporozoites largely lack the ability to perform gliding motility, while the salivary gland sporozoites are highly motile (52). Fig. 8 A shows a schematic of membrane-surface CSP levels with motility. The correlation of low surface CSP levels and low flexibility of MG sporozoites and high CSP-levels and higher flexibility of SG sporozoites (Fig. 8 B) could be a reason for low motility of MG-SPZ *vs* SG-SPZ. However, these are two different developmental stages, differing in several molecular parameters other than CSP, which could contribute to the differential motilities. We measured the CSP levels of 18^th^ day heamocoel, 16^th^, 18^th^ and 19^th^ day salivary gland sporozoites hoping for a significant difference in CSP levels, so that we could make measurements of the Young’s moduli and compare within the SG-stage of the sporozoites. However, the CSP levels were not significantly different for these specific SG-stages (data not shown), and hence a comparison of Young’s moduli for the same stage of parasites was not possible.

**Fig. 8.**
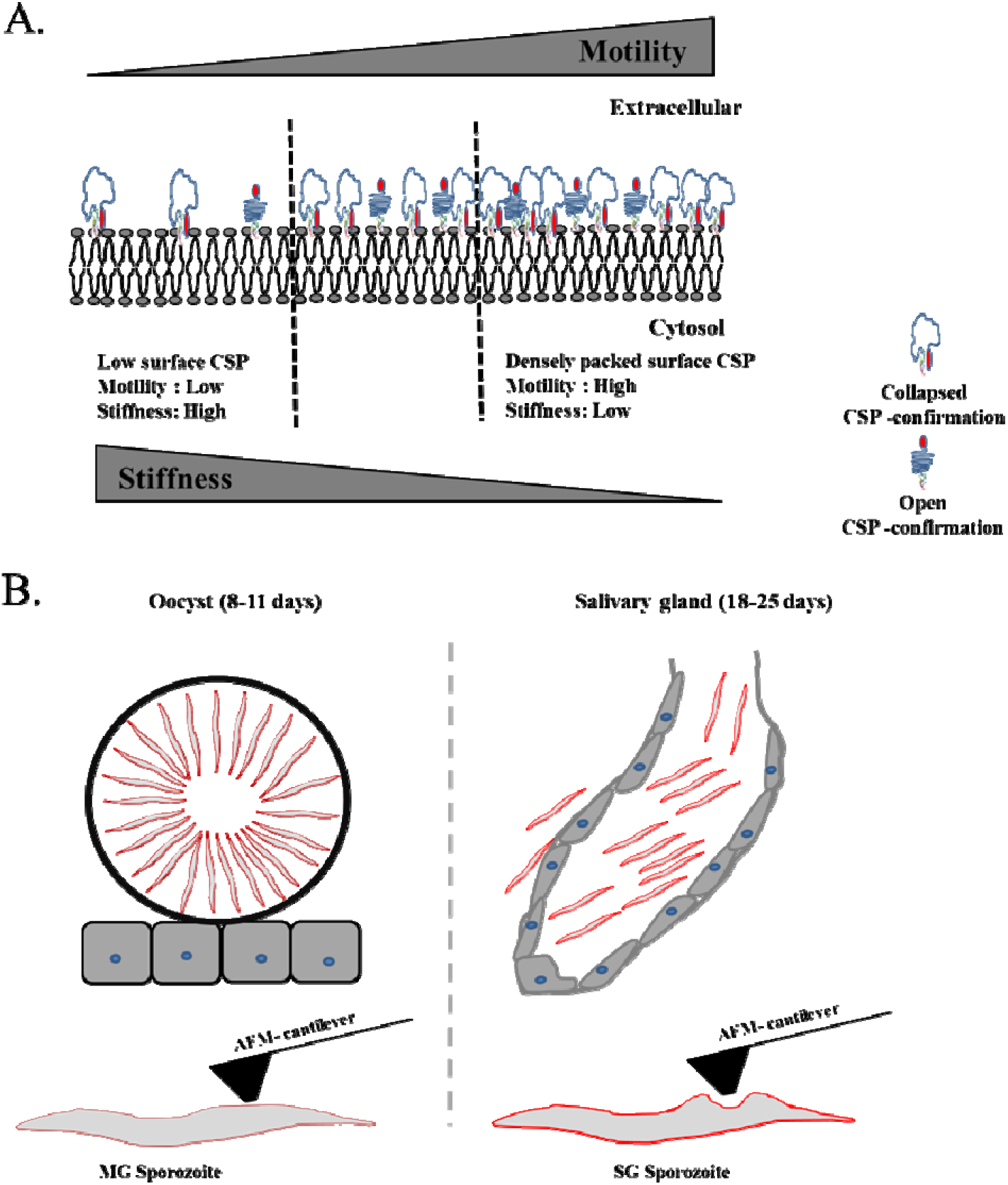
Schematic representation of CSP on sporozoite membrane and its effects on stiffness and motility: A) The quantity of CSP present on membrane inversely regulates membrane stiffness and is directly proportional to its motility. On the cell membrane CSP exists as open and close conformations. B) Changes in surface-CSP levels on sporozoites as they develop from midgut to salivary gland sporozoites.

Experiments with different deletion mutants of CSP producing sporozoites have been difficult because structural integrity of CSP is required for the development of sporozoites (7–10). Electron microscopic studies have shown that most of the CSP mutants, such as those lacking the GPI anchor region (10), some of the 3’-UTR region (8), the repeat region (9) and the N-terminal plus repeat region (9) resulted in deformed sporozoites. However, removal of the one-third of the N-terminal domain of CSP did not appear to interfere with the development of normal-looking oocysts with apparently normal sporozoites within each oocyst (11). This has allowed for comparisons of sporozoite motility of wild-type sporozoites and CSP-Ndel sporozoites (11,12). It was observed that CSP-Ndel sporozoites could invade cell lines with enhanced infectivity, but *in vivo* the sporozoites exhibited problems with migration (11). Such sporozoites had problems exiting the skin infection sites, as also a large prepatent period for development into exoerythrocytic forms in the liver was observed (11). The hypothesis proposed was that the N terminal domain of CSP masked the TSR, and the absence of such a folded conformation impeded the migration of a sporozoite (11).

We studied the motility of CSP-Ndel, CSP-repeat and CSP-NRdel mutants expressed in the *D. discoideum* cells. However, each of these mutants and the irrelevant protein (I27)_4_ exhibited significantly reduced motility as compared to the full-length CSP and exhibited velocity comparable to the control AX2 cells of *D. discoideum* (Fig. 6). This would not have been the case if stiffness was the dominant parameter for cell motility, since CSP-Ndel and CSP-repeat expressing cells exhibited comparable stiffness values to that of the full-length CSP (Fig. 6). Thus, there are other structural/mechanical features that allow the full-length CSP to be the best suited for enhancement of the motility of the cells. The absence of a crystal structure for the full-length CSP remains a lacuna for understanding specific structural features that may contribute towards the motility of sporozoites.

In conclusion, we document for the first time a measurement of the Young’s modulus for *Plasmodium* sporozoites. Using *Dictyostelium* model we also demonstrate that the flexibility and motility of the amoebic cells were significantly enhanced in the presence of the surface-expressed full-length CS-protein. Based on our findings we postulate that the surface expressed CSP is important for *Plasmodium* sporozoites possibly because this protein shields the parasites as they move through different cellular barriers, experiencing considerable friction and drag forces. An analogy may be seen in the free swimming trypanosome bloodstream form, where the abundant variant surface glycoprotein (VSG) provides a shielding function (53,54). Trypanosomes possess a flagellum, and hence the motility is powered quite differently from that of *Plasmodium* sporozoites. However, like CSP, VSGs are also GPI-anchored proteins which adopt two main conformations (4,6,53). The VSGs appear to allow a shielding function which enables a high mobility in the presence of obstacles and confinement (53). It is thus tempting to speculate that the abundant CS-protein on the sporozoite surface may aid the sporozoites perhaps through some mechanical features. An elucidation of the structure of CSP-coated cell-membrane is required to shed further light on the molecular mechanism of contribution of CSP towards enhanced cellular flexibility and motility.

## Materials and Methods

### Plasmids

Both Periv-CSP and Pac15-CSP plasmids were kind gifts from Prof. N. J. Fasel (32). Both plasmids were confirmed by sequencing (Fig. S1). The plasmids were further modified by deleting the N-terminal 18 aa domain and the C-terminal-23 amino acids (putative GPI-anchor) of *PfCSP*, gene leaving 358 amino acids of CSP. Dd *ras* and actin 15 promoter sequences of *D. discoideum* were cloned upstream to *CSP* in Periv-CSP and Pac15-CSP, respectively.

### Anti-CSP Antibodies

The purified recombinant CSP repeat protein ((I_27_)_3_-CSPrep-(I_27_)_3_ (6) was used to generate anti-CSP antibodies in mice. A group of five Balb/c male mice received an intra-peritoneal dose of 100 μg recombinant protein emulsified in Freund’s complete adjuvant. The priming was followed by three booster doses of 50 μg of recombinant protein administered with incomplete Freund’s adjuvant about every 14 days. The polyclonal mouse sera (anti-CSP antibodies) were used in this study along with the commercially available monoclonal antibody anti-(NANP)_5_ (Alpha-diagnostic Cat. No. NANP51-A).

### GPI-Phospolipase D (PLD)

Nearly 3 units of GPI-PLD enzyme (Sigma Cat. no. SML0566) were used for the experiments. The *D. discoideum* cells were incubated with the GPI-PLD enzyme for different time point, washed thrice with fresh culture media, harvested and used for flow cytometry and cell-motility assays.

### *D. discoideum* cell culture and transfection

The *D. discoideum* cells (AX2) were grown in HL5 media at 22°C in the presence of 1mg/ml Penstrep (Invitrogen) and with appropriate antibiotic drug in 10 cm petriplate as described earlier (32). *D. discoideum* cells were transfected using electroporation as described previously (55). For induction of the CSP-protein, cells were washed, appropriate amounts of cAMP were added for 60 minutes and washed again prior to further experimentation.

### Flow cytometry assays

Flow cytometry assays were carried out using nearly ~10^7^ *D. discoideum* cells. The cells were centrifuged at 1000 *g* for 5 min and re-suspended in 200 μl of PBS containing 3% BSA. The cells were fixed in 4% paraformaldehyde for 15 min followed by blocking in 5% BSA for 1 hr. The cells were stained with anti-(NANP)_5_ antibodies (1:500 dilution) for 1 hr. Cells were further incubated for 1 hr in solution containing goat anti-rabbit antibody conjugated with FITC (BD Biosciences, USA). All steps were carried out at room temperature. The cells were washed and acquired on BD LSR Fortessa and the mean fluorescent intensity (MFI) data was determined using FACS Diva software.

### Cell indentation assays using AFM

*D. discoideum* cells (~3×10^4^) were grown overnight in 35 mm dishes containing the HL5 media. The medium was then removed, and the adherent cells were washed gently with PBS buffer and subsequently submerged in PBS buffer (pH 7.4) and assayed within 1 hr at room temperature. For the cAMP induction, cells were incubated with different concentrations of cAMP for 60 min, washed and then used for AFM studies. AFM (Nanowizard-3 model) from JPK Instruments (Germany) was used for all cell indentation assays. Force vs distance curves were obtained at a pulling speed of 400nm/s. ‘V-shaped’ cantilevers with conical-shaped silicon-nitride tips from Bruker (USA) were used for indentation assays. Cantilevers were calibrated prior to indentation measurements based on equipartition theorem. The spring constant of the cantilevers was measured to be ~35 pN/nm. Two or more indentions were performed on each cell and the mean values of Young’s modulus of each cell were plotted in the dot plots.

### Data Analysis

Young’s modulus was calculated from the force vs distance cell indentation curves by using the Hertz model of elasticity for a conical indenter (56)(56). The indentation region of each force vs distance trace was fitted to the following equation to extract Young’s modulus (E):

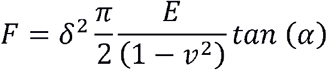

Where F, υ, α, and δ denote force, Poisson’s ratio, open cone angle of the cantilever-tip and depth of indentation, respectively. The indentation region of the force vs distance traces was used to fit the Hertz model equation shown above to measure the Young’s modulus.

### Cell motility assays

Motility of *D. discoideum* cells was measured using the chemoattractant folate in 1% agarose as described in Laevsky et al. (35). *D. discoideum c*ells (~3×10^3^) were loaded in one trough, when the other trough had been preincubated for 1 hr with 0.1 mM folate. The motility was monitored at 20x magnification, through 1% agarose-gel by taking time lapse video with 5 sec frames for about 8-10 min. The path of each cell movement was tracked by manual tracking using ImageJ software.

### Generation and isolation of *P. berghei* sporozoites

Swiss albino mice were infected with *Plasmodium berghei* ANKA parasites. After obtaining 2-3% parasitemia, the blood was collected in anticoagulant and diluted to 1% parasitemia. Nearly 200 μl of the diluted blood was injected intraperitoneally into each mouse. On day 3 post infection, blood smears were made, and Giemsa stained to check for the presence of gametocytes. All gametocyte positive mice were anesthetized and placed on top of a cage harboring female *Anopheles stephensi* mosquitoes. The blood meal was given to mosquitoes for a duration of 15 minutes, for 2 successive days. The blood fed mosquitoes were maintained in an environmental chamber at 21°C and 80% RH. On days 14 and 18-20 post infection respectively, the mosquito midguts and salivary glands were dissected and placed in 1.5 ml Eppendorf tube containing incomplete DMEM. Both tissues were mechanically disrupted using a plastic pestle and centrifuged at 800 RPM for 3 min at 4°C. The supernatant containing partially purified sporozoites were used for performing IFA, Western blotting, flow cytometer and AFM assays.

### Sporozoite IFA

Both oocyst and salivary gland sporozoites were spotted on a clean glass slide. Sporozoites were fixed with 4% formalin solution (SIGMA, Cat# HT50-1-2) for 20 min at room temperature. Sporozoites were washed with 1XTBS followed by permeabilization with acetone:methanol solution used in 1:3 ratio. Nonspecific blocking was done with 3% BSA in 1XTBS for 1 h at 37°C. Following nonspecific blocking, sporozoites were stained with mouse 3D11 monoclonal antibody (57), (1:1000 dilution in 1% BSA) that recognizes the repeats of *P. berghei* CSP protein. The immunoreactivity was revealed by using goat anti-mouse secondary antibody conjugated to Alexa Fluor 594 (1:300 dilution in 1% BSA) (Molecular probes, Cat# A-11005). The CSP immunostaining was visualized under fluorescence microscope. Fluorescence intensities were measured using AR software for both midgut and salivary gland sporozoites (25 sporozoites from each group) and plotted in dot plot graph.

### Western Blotting

Equal number of sporozoites (3×10^4^) were lysed in SDS sample buffer by boiling at 100°C for 8 min. Lysates were resolved on 10% SDS-PAGE and electrophoretically transferred to nitrocellulose membrane. Nonspecific blocking of nitrocellulose membrane was done with 3% skimmed milk powder in 1XTBS. Following nonspecific blocking, nitrocellulose membrane was incubated with mouse 3D11 antibody (1:1000 dilution in 1XTBS) for 1 h at 37°C, followed by washes and subsequent incubation with anti-mouse secondary antibody conjugated to horseradish peroxidase (1:5000 dilution in 1XTBS) (Bangalore Genei, Cat# 105502) for 1 h at 37°C. The membrane was developed by using ECL prime western blotting detection kit (Amersham) and images were captured using Versadoc (Biorad).

### AFM assay of sporozoites

Midgut and salivary gland sporozoites were dissected and harvested in 3% BSA and DMEM solution and centrifuged at 1000 *g* for 5 min at 4°C. About 20,000 sporozoites/100 μl was spotted on the polylysine-coated cover slide. Sporozoites were visualized with a 20x magnified lens of the microscope over which AFM was mounted. Isolated sporozoites were identified and they were indented around the middle for the stiffness measurements. All AFM measurements were made within 40 min of mosquito dissection. The sporozoites were alive during the AFM measurements as determined by their twitching movements. Multiple measurements (at least 2) were measured for each sporozoite, and the data was averaged/cell for the dot plot. A total of 17 MG and 18 SG sporozoites were probed using AFM.

## Acknowledgements

We thank Prof. N. Fasel from (Department of Biochemistry, University of Lausanne, Basel Switzerland) for the kind gifts of Periv CSP and Pac15 CSP plasmid constructs, Prof. Roop Mallik and Mr. Ashwin D’souza from TIFR for their guidance with culturing and video microscopy of *D. discoideum* cells. The authors acknowledge financial support from TIFR-DAE. Funding for Nikon AR fluorescent microscope from DBT (Grant/Award number: BT/PR2495/BRB/10/950/2011) to Prof. Kota Arun Kumar, Department of Animal Sciences, University of Hyderabad is also acknowledged.

## Author contributions

AP, CD, VP, conception and design, acquisition of data, analysis and interpretation of data, drafting the article; AP, SRR, KAK, DS, generation and experiments with *Plasmodium* sporozoites; SRKA, SS, conception and design, analysis and interpretation of data, drafting the article, and guiding the study.

