## Supplemental document for "Surface expressed *Plasmodium* circumsporozoite protein (CSP) modulates cellular flexibility and motility"

Shorter Title (for Mobile devices and RSS feeds): *Plasmodium* CSP modulates cellular flexibility and motility

**Aditya Prasad Patra[a], Vrushali Pathak[a], Segireddy Rameswara Reddy[b] , Aditya Chhatre[a], Crismita Dmello[a], Satya Narayan[c], Dipti Singh[b], Kota Arun Kumar[b], Sri Rama Koti Ainavarapu[c,1] and Shobhona Sharma[a,1]**

A.

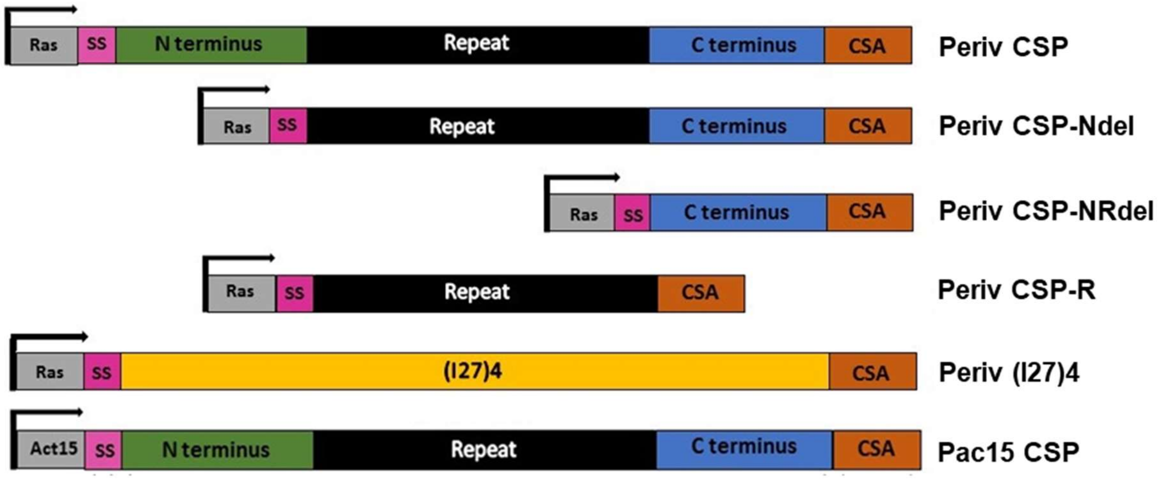


B.

### Periv CSP

------------------------------------------------------------------------TAGGGAGACCG
GAATTCGAGCTCGCCC**CATAATATTTTTGTGTTCTTATAATTTGGTTAAATCGATGAATAATATTTGATTAGTATATGTTTTTTTTTCCTT**
**TTTTTTATTTTTATTTTTATTTTTTTAAAAAATAAAAATTAGAATAAATATTTCTATTTGAAGGAGTTTTTATTTGTATTTAAAATTATAT**
**TAAACATAGTGAACCTAAAAATAGATTTGTGACGGTATATGATAAGAAAATTCTAAAAAAAAAATTCAGATAATTTTTGGATTGGAAACAA**
**CAACCAAAAAAAAAAAAAAAAAAAAAAAAAAAAAAATCAAAAAAAAAAAAAAAAAAAAAATTAAAATCAAAAAAAAAAGGTATTTAAAGAAATT**
**TTTTAAAATATTATTATATATCTTTAATTGGGCAAAACACACTTTTAACACGC**CCAAGCCGGGATAAAATCATTAAATTGAAAAATTAAAA
TTTCATACAAATTATCTTTTTAAATAATAAAATTTATAAAATGTCTAGATTTTTAGTATTGATAATATTATATAATATTTTAAATAGTGCA
CATTCAGCTCCAACCCAGGATCCCCGGGTACCATTATTCCAGGAATACCAGTGCTATGGAAGTTCGTCAAACACAAGGGTTCTAAATGAAT
TAAATTATGATAATGCAGGCACTAATTTATATAATGAATTAGAAATGAATTATTATGGGAAACAGGAAAATTGGTATAGTCTTAAAAAAAG
TAGTAGATCACTTGGAGAAAATGATGATGGAAATAACGAAGACAACGAGAAATTAAGGAAACCAAAACATAAAAAATTAAAGCAACCAGCG
GATGGTAATCCTGATCCAAATGCAAACCCAAATGTAGATCCCAATGCCAACCCAAATGTAGATCCAAATGCAAACCCAAATGTAGATCCAA
ATGCAAACCCAAATGCAAACCCAAATGCAAACCCAAATGCAAACCCAAATGCAAACCCAAATGCAAACCCAAATGCAAACCCAAATGCAAA
CCCAAATGCAAACCCAAATGCAAACCCAAATGCAAACCCAAATGCAAACCCAAATGCAAACCCAAATGCAAACCCCAATGCAAATCCTAAT
GCAAACCCAAATGCAAACCCAAACGTAGATCCTAATGCAAATCCAAATGCAAACCCAAACGCAAACCCCAATGCAAATCCTAATGCAAACC
CCAATGCAAATCCTAATGCAAATCCTAATGCCAATCCAAATGCAAATCCAAATGCAAACCCAAACGCAAACCCCAATGCAAATCCTAATGC
CAATCCAAATGCAAATCCAAATGCAAACCCAAATGCAAACCCAAATGCAAACCCCAATGCAAATCCTAATAAAAACAATCAAGGTAATGGA
CAAGGTCACAATATGCCAAATGACCCAAACCGAAATGTAGATGAAAATGCTAATGCCAACAGTGCTGTAAAAAATAATAATAACGAAGAAC
CAAGTGATAAGCACATAAAAGAATATTTAAACAAAATACAAAATTCTCTTTCAACTGAATGGTCCCCATGTAGTGTAACTTGTGGAAATGG
TATTCAAGTTAGAATAAAGCCTGGCTCTGCTAATAAACCTAAAGACGAATTAGATTATGCAAATGATATTGAAAAAAAAATTTGTAAAATG
GAAAAATGTGGATCTCCAACTCCAACTGAAACAGCCACCCCATCTCCAACTCCAACTGAAACAGCCACCCCATCTCCAACTCCAAAACCAA
CCAGCACACCAGAAGAAACTGAAGCACCTTCATCAGCAACAACTCTTATTTCACCATTATCTTTAATTGTTATTTTCATTTCTTTTGTTTT
ATTAATTTAACTCGAG

### Periv CSP-Ndel

ATGTCTAGATTTTTAGTATTGATAATATTATATAATATTTTAAATAGTGCACATTCAGCTCCAACCCAGGATCCCCGGGTACCAAATCCTG
ATCCAAATGCAAACCCAAATGTAGATCCCAATGCCAACCCAAATGTAGATCCAAATGCAAACCCAAATGTAGATCCAAATGCAAACCCAAA
TGCAAACCCAAATGCAAACCCAAATGCAAACCCAAATGCAAACCCAAATGCAAACCCAAATGCAAACCCAAATGCAAACCCAAATGCAAAC
CCAAATGCAAACCCAAATGCAAACCCAAATGCAAACCCAAATGCAAACCCAAATGCAAACCCCAATGCAAATCCTAATGCAAACCCAAATG
CAAACCCAAACGTAGATCCTAATGCAAATCCAAATGCAAACCCAAACGCAAACCCCAATGCAAATCCTAATGCAAACCCCAATGCAAATCC
TAATGCAAATCCTAATGCCAATCCAAATGCAAATCCAAATGCAAACCCAAACGCAAACCCCAATGCAAATCCTAATGCCAATCCAAATGCA
AATCCAAATGCAAACCCAAATGCAAACCCAAATGCAAACCCCAATGCAAATCCTAATAAAAACAATCAAGGTAATGGACAAGGTCACAATA
TGCCAAATGACCCAAACCGAAATGTAGATGAAAATGCTAATGCCAACAGTGCTGTAAAAAATAATAATAACGAAGAACCAAGTGATAAGCA
CATAAAAGAATATTTAAACAAAATACAAAATTCTCTTTCAACTGAATGGTCCCCATGTAGTGTAACTTGTGGAAATGGTATTCAAGTTAGA
ATAAAGCCTGGCTCTGCTAATAAACCTAAAGACGAATTAGATTATGCAAATGATATTGAAAAAAAAATTTGTAAAATGGAAAAATGTGGAT

CTCCAACTCCAACTGAAACAGCCACCCCATCTCCAACTCCAACTGAAACAGCCACCCCATCTCCAACTCCAAAACCAACCAGCACACCAGA
AGAAACTGAAGCACCTTCATCAGCAACAACTCTTATTTCACCATTATCTTTAATTGTTATTTTCATTTCTTTTGTTTTATTAATTTAACTC
GAG

### Periv CSP-NRdel

**ATG**TCTAGATTTTTAGTATTGATAATATTATATAATATTTTAAATAGTGCACATTCAGCTCCAACCCAGGATCCCCGGGTACCAAATATGC
CAAATGACCCAAACCGAAATGTAGATGAAAATGCTAATGCCAACAGTGCTGTAAAAAATAATAATAACGAAGAACCAAGTGATAAGCACAT
AAAAGAATATTTAAACAAAATACAAAATTCTCTTTCAACTGAATGGTCCCCATGTAGTGTAACTTGTGGAAATGGTATTCAAGTTAGAATA
AAGCCTGGCTCTGCTAATAAACCTAAAGACGAATTAGATTATGCAAATGATATTGAAAAAAAAATTTGTAAAATGGAAAAATGTGGATCTC
CAACTCCAACTGAAACAGCCACCCCATCTCCAACTCCAACTGAAACAGCCACCCCATCTCCAACTCCAAAACCAACCAGCACACCAGAAGA
AACTGAAGCACCTTCATCAGCAACAACTCTTATTTCACCATTATCTTTAATTGTTATTTTCATTTCTTTTGTTTTATTAATTTAACTCGAG

### Periv CSP-R

**ATG**TCTAGATTTTTAGTATTGATAATATTATATAATATTTTAAATAGTGCACATTCAGCTCCAACCCAGGATCCCCGGGTACCAAATCCTG
ATCCAAATGCAAACCCAAATGTAGATCCCAATGCCAACCCAAATGTAGATCCAAATGCAAACCCAAATGTAGATCCAAATGCAAACCCAAA
TGCAAACCCAAATGCAAACCCAAATGCAAACCCAAATGCAAACCCAAATGCAAACCCAAATGCAAACCCAAATGCAAACCCAAATGCAAAC
CCAAATGCAAACCCAAATGCAAACCCAAATGCAAACCCAAATGCAAACCCAAATGCAAACCCCAATGCAAATCCTAATGCAAACCCAAATG
CAAACCCAAACGTAGATCCTAATGCAAATCCAAATGCAAACCCAAACGCAAACCCCAATGCAAATCCTAATGCAAACCCCAATGCAAATCC
TAATGCAAATCCTAATGCCAATCCAAATGCAAATCCAAATGCAAACCCAAACGCAAACCCCAATGCAAATCCTAATGCCAATCCAAATGCA
AATCCAAATGCAAACCCAAATGCAAACCCAAATGCAAACCCCAATGCAAATCCTAATAAAAACAATCAAGGTAATGGACAAGGTCACGGAT
CTCCAACTCCAACTGAAACAGCCACCCCATCTCCAACTCCAACTGAAACAGCCACCCCATCTCCAACTCCAAAACCAACCAGCACACCAGA
AGAAACTGAAGCACCTTCATCAGCAACAACTCTTATTTCACCATTATCTTTAATTGTTATTTTCATTTCTTTTGTTTTATTAATTTAACTC
GAG

### Periv (I27)4

**ATG**TCTAGATTTTTAGTATTGATAATATTATATAATATTTTAAATAGTGCACATTCAGCTCCAACCCAGGATCCCCGGGTACCACATCATC
ACCATCATCATGGATCCCTAATAGAAGTGGAAAAGCCTCTGTACGGAGTAGAGGTGTTTGTTGGTGAAACAGCCCACTTTGAAATTGAACT
TTCTGAACCTGATGTTCACGGCCAGTGGAAGCTGAAAGGACAGCCTTTGGCAGCTTCCCCTGACTGTGAAATCATTGAGGATGGAAAGAAG
CATATTCTGATCCTTCATAACTGTCAGCTGGGTATGACAGGAGAGGTTTCCTTCCAGGCTGCTAATACCAAATCTGCAGCCAATCTGAAAG
TGAAAGAATTGAGATCCCTAATAGAAGTGGAAAAGCCTCTGTACGGAGTAGAGGTGTTTGTTGGTGAAACAGCCCACTTTGAAATTGAACT
TTCTGAACCTGATGTTCACGGCCAGTGGAAGCTGAAAGGACAGCCTTTGGCAGCTTCCCCTGACTGTGAAATCATTGAGGATGGAAAGAAG
CATATTCTGATCCTTCATAACTGTCAGCTGGGTATGACAGGAGAGGTTTCCTTCCAGGCTGCTAATACCAAATCTGCAGCCAATCTGAAAG
TGAAAGAATTGAGATCCCTAATAGAAGTGGAAAAGCCTCTGTACGGAGTAGAGGTGTTTGTTGGTGAAACAGCCCACTTTGAAATTGAACT
TTCTGAACCTGATGTTCACGGCCAGTGGAAGCTGAAAGGACAGCCTTTGGCAGCTTCCCCTGACTGTGAAATCATTGAGGATGGAAAGAAG
CATATTCTGATCCTTCATAACTGTCAGCTGGGTATGACAGGAGAGGTTTCCTTCCAGGCTGCTAATACCAAATCTGCAGCCAATCTGAAAG
TGAAAGAATTGAGATCCGAGCTCCTAATAGAAGTGGAAAAGCCTCTGTACGGAGTAGAGGTGTTTGTTGGTGAAACAGCCCACTTTGAAAT
TGAACTTTCTGAACCTGATGTTCACGGCCAGTGGAAGCTGAAAGGACAGCCTTTGGCAGCTTCCCCTGACTGTGAAATCATTGAGGATGGA
AAGAAGCATATTCTGATCCTTCATAACTGTCAGCTGGGTATGACAGGAGAGGTTTCCTTCCAGGCTGCTAATACCAAATCTGCAGCCAATC
TGAAAGTGAAAGAATTGGCTAGCAGATCTTGTTGCGGATCTCCAACTCCAACTGAAACAGCCACCCCATCTCCAACTCCAACTGAAACAGC
CACCCCATCTCCAACTCCAAAACCAACCAGCACACCAGAAGAAACTGAAGCACCTTCATCAGCAACAACTCTTATTTCACCATTATCTTTA
ATTGTTATTTTCATTTCTTTTGTTTTATTAATTTAACTCGAG

# C.

### Pac15 CSP

----------------------------------------------------------TAGGGAGACCGGAATTCGAGCTCGCCC**ACAAAT**
**TAATTAATCCCATCAAATCTTTAAAAAAAAAAATGGTTTAAAAAAACTTGGGTTGGTTAATTATTATTTGAAAATTTTAAAACCCAAATTA**
**AAAAAAAAAATGGGATTCAAAAATTTTTTTTTTTTTTTTTTTTTTTTTTTTTTTTTTTTTTTTTTCAAATTGGAAAAAAAAATTTTTTTTTTT**
**TTTTTTTTTTTTTTTAAAAAAAAAAATTTATTTTAAAAAAAAAAAAAAATGGA**CCAAGCCGGGATAAAATCATTAAATTGAAAAATTAAAA
TTTCATACAAATTATCTTTTTAAATAATAAAATTTATAAA**ATG**TCTAGATTTTTAGTATTGATAATATTATATAATATTTTAAATAGTGCA
CATTCAGCTCCAACCCAGGATCCCCGGGTACCATTATTCCAGGAATACCAGTGCTATGGAAGTTCGTCAAACACAAGGGTTCTAAATGAAT

TAAATTATGATAATGCAGGCACTAATTTATATAATGAATTAGAAATGAATTATTATGGGAAACAGGAAAATTGGTATAGTCTTAAAAAAAG
TAGTAGATCACTTGGAGAAAATGATGATGGAAATAACGAAGACAACGAGAAATTAAGGAAACCAAAACATAAAAAATTAAAGCAACCAGCG
GATGGTAATCCTGATCCAAATGCAAACCCAAATGTAGATCCCAATGCCAACCCAAATGTAGATCCAAATGCAAACCCAAATGTAGATCCAA
ATGCAAACCCAAATGCAAACCCAAATGCAAACCCAAATGCAAACCCAAATGCAAACCCAAATGCAAACCCAAATGCAAACCCAAATGCAAA
CCCAAATGCAAACCCAAATGCAAACCCAAATGCAAACCCAAATGCAAACCCAAATGCAAACCCAAATGCAAACCCCAATGCAAATCCTAAT
GCAAACCCAAATGCAAACCCAAACGTAGATCCTAATGCAAATCCAAATGCAAACCCAAACGCAAACCCCAATGCAAATCCTAATGCAAACC
CCAATGCAAATCCTAATGCAAATCCTAATGCCAATCCAAATGCAAATCCAAATGCAAACCCAAACGCAAACCCCAATGCAAATCCTAATGC
CAATCCAAATGCAAATCCAAATGCAAACCCAAATGCAAACCCAAATGCAAACCCCAATGCAAATCCTAATAAAAACAATCAAGGTAATGGA
CAAGGTCACAATATGCCAAATGACCCAAACCGAAATGTAGATGAAAATGCTAATGCCAACAGTGCTGTAAAAAATAATAATAACGAAGAAC
CAAGTGATAAGCACATAAAAGAATATTTAAACAAAATACAAAATTCTCTTTCAACTGAATGGTCCCCATGTAGTGTAACTTGTGGAAATGG
TATTCAAGTTAGAATAAAGCCTGGCTCTGCTAATAAACCTAAAGACGAATTAGATTATGCAAATGATATTGAAAAAAAAATTTGTAAAATG
GAAAAATGTGGATCTCCAACTCCAACTGAAACAGCCACCCCATCTCCAACTCCAACTGAAACAGCCACCCCATCTCCAACTCCAAAACCAA
CCAGCACACCAGAAGAAACTGAAGCACCTTCATCAGCAACAACTCTTATTTCACCATTATCTTTAATTGTTATTTTCATTTCTTTTGTTTT
ATTAATTTAACTCGAG

**Fig. S1. Schematic drawing and nucleotide sequences of the constructs used. A**) Cartoon of Periv-CSP and Pac15-CSP constructs. The signal and anchor regions of *PfCSP* gene were removed and replaced with those of the *D. discoideum* contact surface antigen (*CSA)* gene leaving 358 amino acids of CSP. **B)** and **C)** Panels show the flanking nucleotide sequences of all Periv-CSP constructs and Pac15-CSP, respectively. Ras and actin15 promoter sequences of *D. discoideum* were used in the Periv-CSP and Pac15-CSP constructs, respectively (shown as black, bold and underlined). The signal (pink) and the GPI-anchor (brown) of the *CSA* gene of *D. discoideum* was used in both the constructs. The ATG (blue) and N terminal region (green), repeat region (black), C terminal region (light blue) of the *P. falciparum* CSP gene sequence are marked in respective colors.

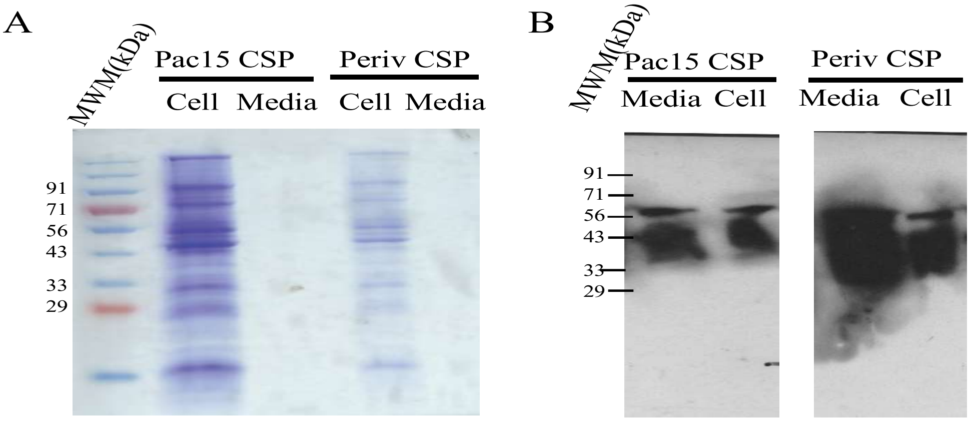


**Fig. S2.** Coomassie-stained gel (**A**) and immunoblots (**B**) of the expressed proteins from total cell lysate and those secreted in the culture media. Anti-PfCSP antibodies were used as the probe in immunoblots.

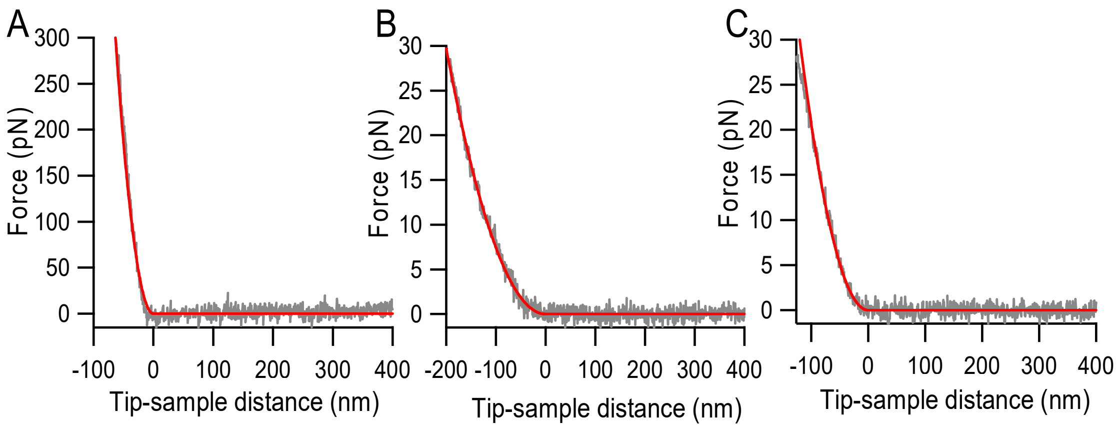


**Fig. S3.** Representative force vs distance traces of AX2 cells (A), Periv-CSP (B) and Pac15-CSP (C) obtained in AFM indentation assays.

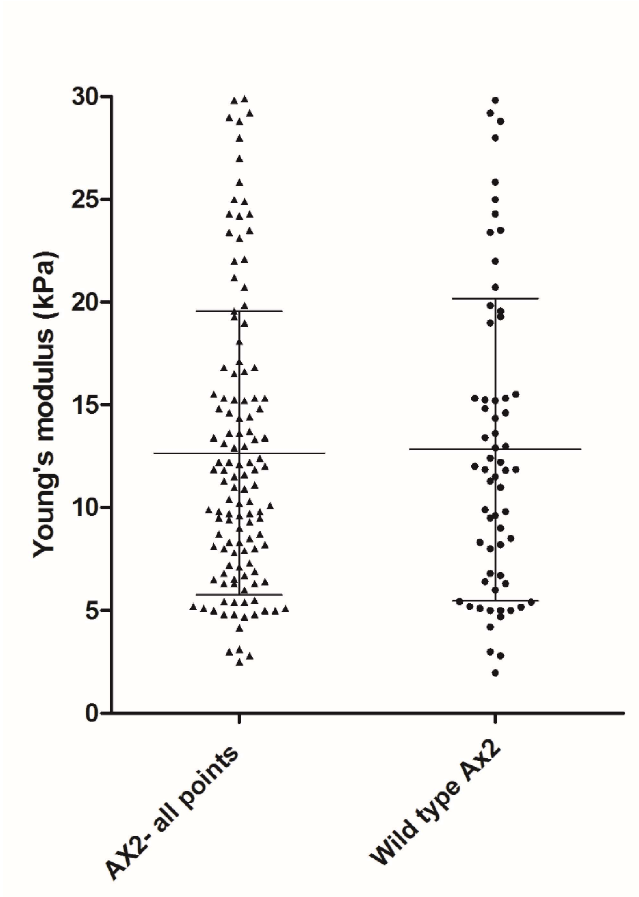


**Fig. S4.** Dot plot of the measured stiffness of the AX2 cells. Two points were measured on each AX2 cell and the mean of these two points is calculated. In the plot the first column shows individual indentation points on the AX2 control cell and second column "Wild type AX2" shows the mean values of measured points per cell. No statistically significant difference was observed between these two analyses.

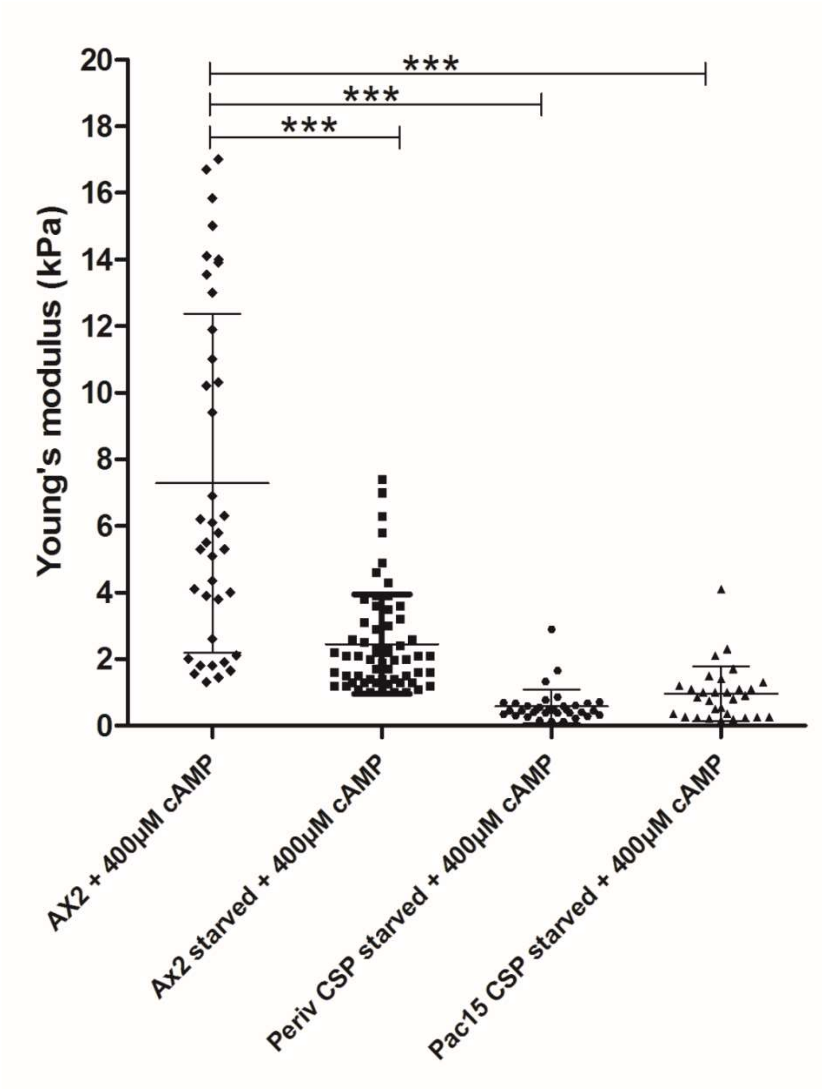


**Fig. S5.** Dot plot of measured stiffness of various *D. discoideum* cells under starved condition in the presence of 400 uM cAMP. ***$p$<0.001.

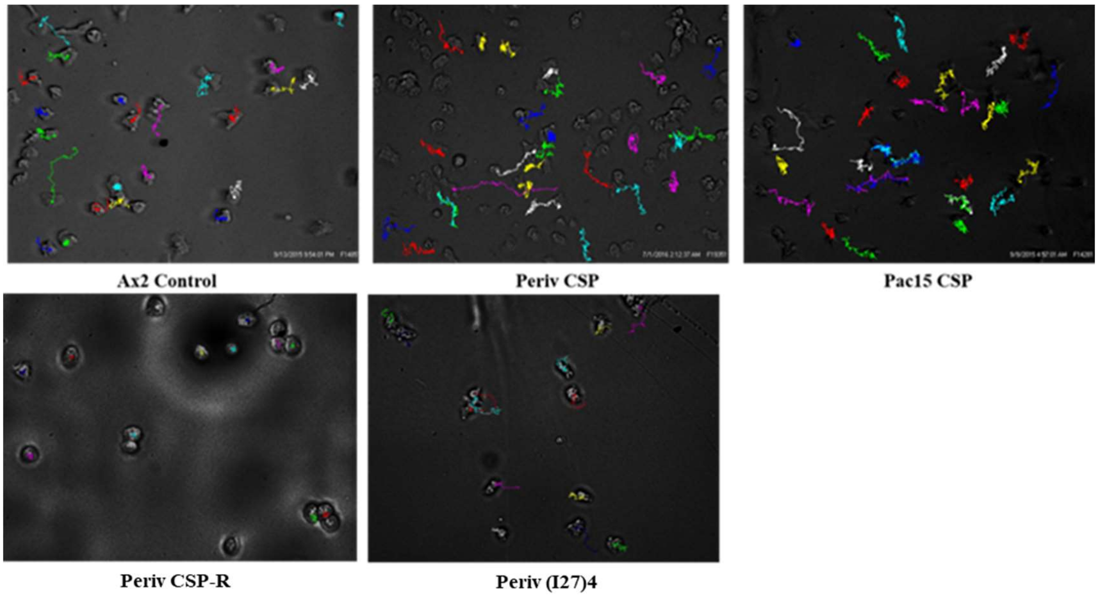


**Fig. S6.** ***D. discoideum*** **cell motility tracks.** The cell motility was measured in 1% agarose-gel by taking time lapse video with 5 sec frames. The color path of each cell movement was tracked by manual tracking using ImageJ software. Here we provide the final snapshot images of representative video tracks after 8 min for AX2, Periv CSP, Pac15 CSP, Periv CSP-R and Periv (I27)4 transfected *Dictyostelium* cells induced with 400 $\mu$M cAMP**.**

**A.**

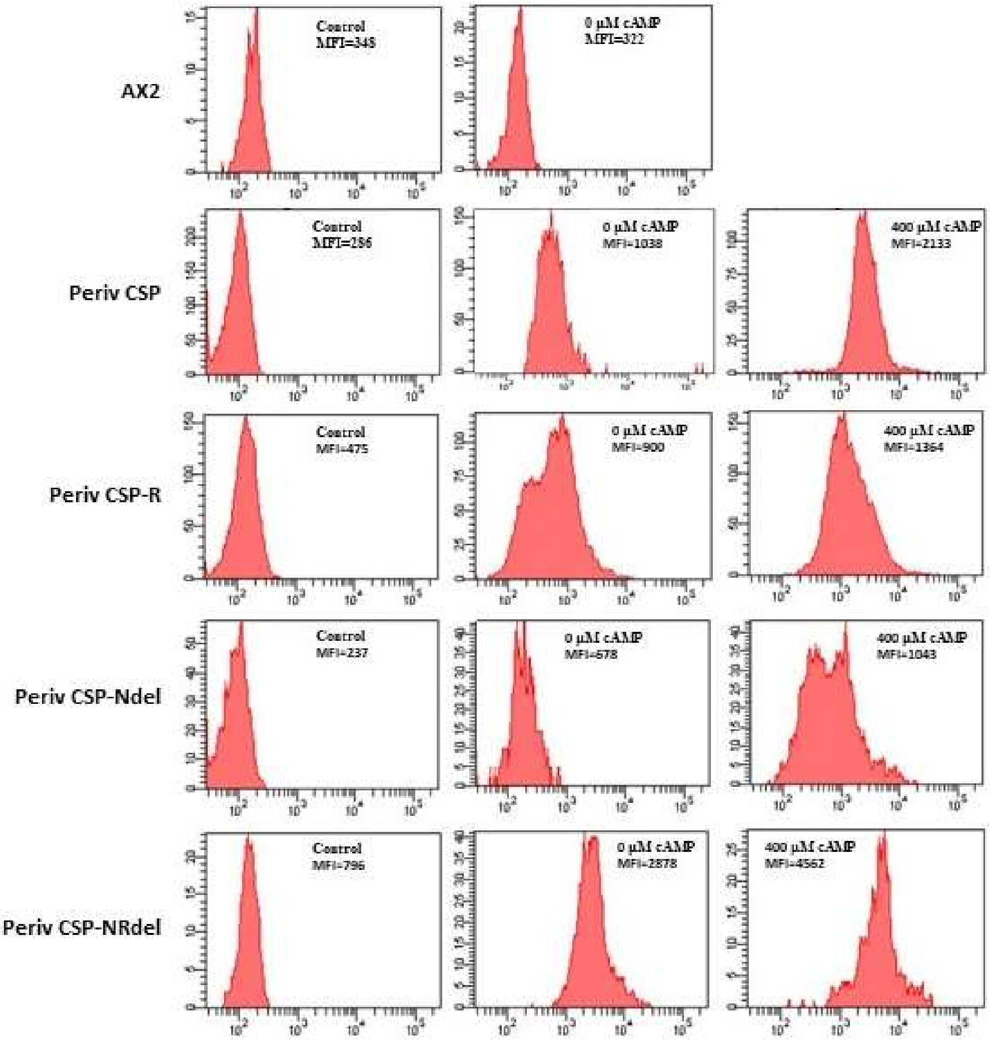


**B.**

| **cAMP (µM)** | **Mean fluorescent intensity (MFI)** | | | |
|---|---|---|---|---|
| | Periv CSP | Periv CSP-R | Periv CSP-Ndel | Periv CSP-NRdel |
| 0 | 852 ± 185 | 857 ± 43 | 725 ± 47 | 2806 ±72 |
| 400 | 2358 ± 225 | 1469 ± 105 | 1137 ± 94 | 4464 ± 79 |

**Fig S7.** A) Representative images of mean fluorescence intensity (MFI) of surface protein expression on AX2, Periv CSP, Periv CSP-R, Periv CSP-Ndel and Periv CSP-NRdel transfected *D. discoideum* after incubation with 0 µM and 400 µM cAMP. The vertical panels left to right show secondary antibody control, 0 µM and 400 µM cAMP treated cells for each construct. B) The measured MFI values shown are the average ± SD of surfaced expressed CSP.

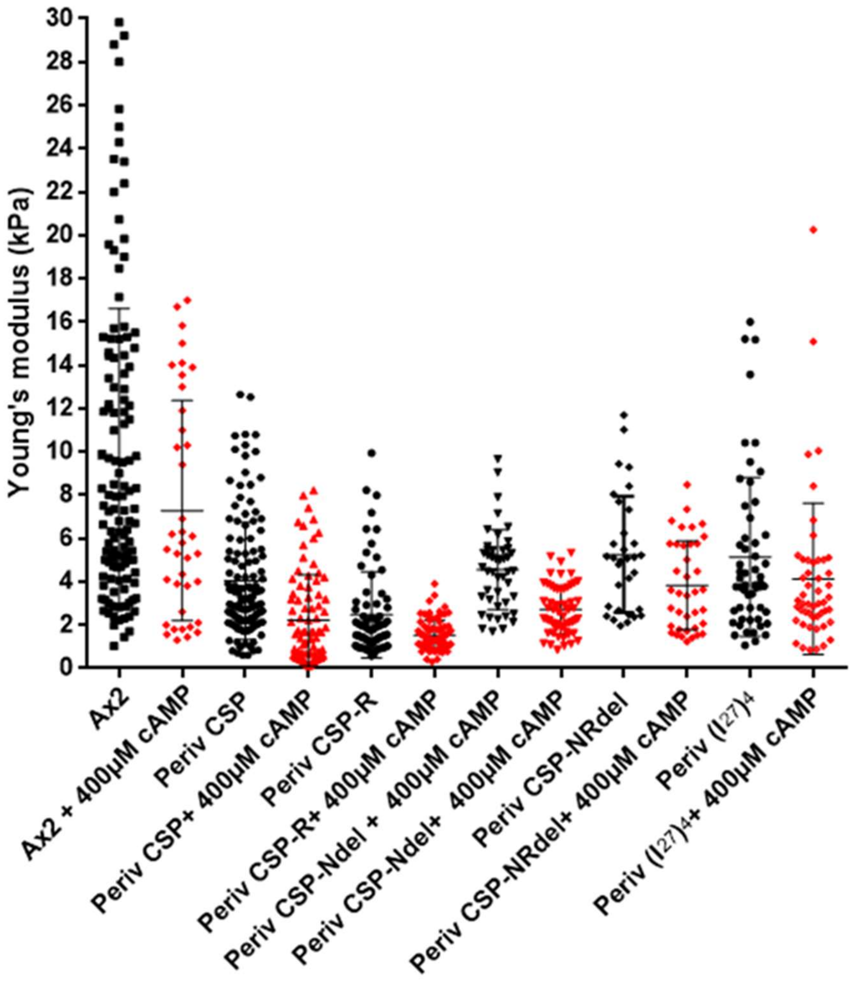


**Fig. S8.** Dot plot of measured stiffness of various constructs of CSP and (I27)4 expressed in *D. discoideum* cells in the presence of 0 µM and 400 µM cAMP.

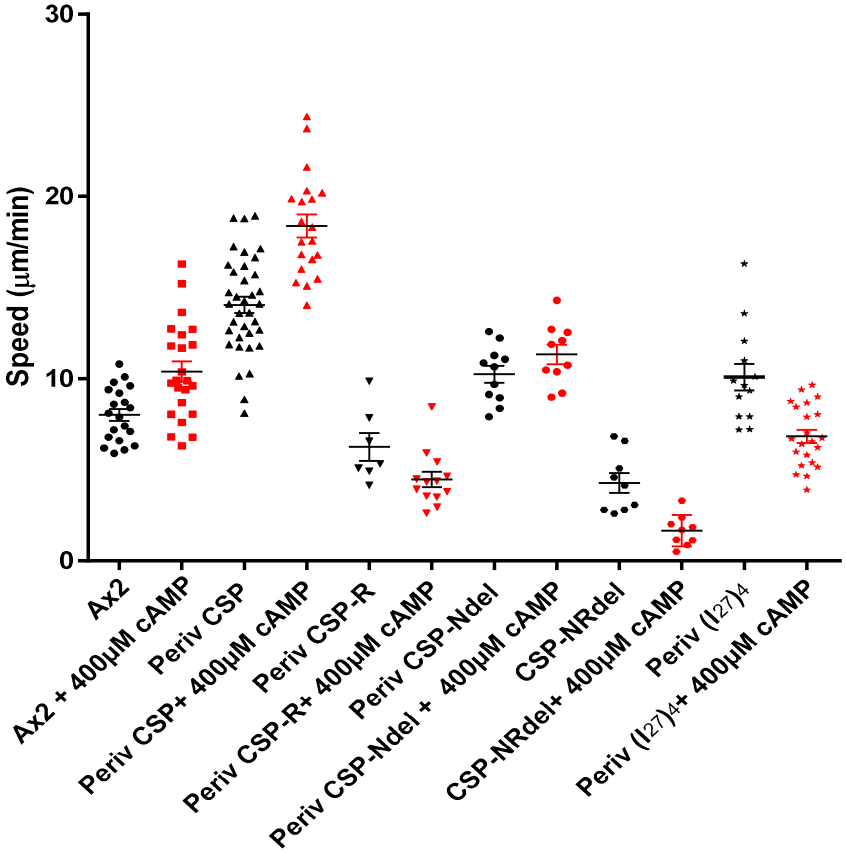


**Fig. S9.** Dot plot of speed of various constructs of CSP and (I27)4 expressed in *D. discoideum* cells through agar plate in the presence of 0 $\mu$M and 400 $\mu$M cAMP.

| cAMP (µM) | Mean fluorescent intensity (MFI) | | Young's modulus (kPa) | |
|---|---|---|---|---|
| | Periv-CSP | Pac15-CSP | Periv-CSP | Pac15-CSP |
| 0 | 321 ± 42 | 275 ± 25 | 3.6 ± 2.4 | 3.2 ± 3.5 |
| 50 | 629 ± 23 | 573 ± 29 | 2.0 ± 1.8 | 2.5 ± 2.9 |
| 100 | 806 ± 65 | 862 ± 37 | 1.7 ± 1.0 | 2.1 ± 1.9 |
| 400 | 2824 ± 47 | 1335 ± 29 | 1.2 ± 1.6 | 1.0 ± 0.9 |

**Table S1.** cAMP dependent surface-CSP expression levels measured by flow cytometer assays and the corresponding cell stiffness measured by AFM indentation assays for Periv-CSP and Pac15-CSP transfected *D. discoideum* cells. The data shown are average ± SD.

| Construct | Speed (µm/min) | | Young's modulus (kPa) | |
|---|---|---|---|---|
| | 0 µM cAMP | 400 µM cAMP | 0 µM cAMP | 400 µM cAMP |
| $AX_2$ | 8.1 ± 1.5 | 10.4 ± 2.6 | 9.7 ± 6.9 | 7.3 ± 5.1 |
| Per iv CSP | 14.4 ± 2.7 | 18.5 ± 2.8 | 4 ± 2.7 | 2.2 ± 2.1 |
| Pac15 CSP | 11.9 ± 2.1 | 19.5 ± 3.8 | 3.2 ± 3.5 | 0.95 ± 0.8 |
| Periv CSP-R | 6.3 ± 2 | 4.5 ± 1.5 | 2.5 ± 2 | 1.5 ± 0.7 |
| Periv CSP-Ndel | 10.2 ± 1.5 | 11.3 ± 1.6 | 4.5 ± 1.8 | 2.7 ± 1.1 |
| Periv CSP-NRdel | 4.8 ± 1.6 | 1.7 ± 0.8 | 5.3 ± 2.6 | 3.8 ± 2.1 |
| (I27)4 | 10.1 ± 1.6 | 6.8 ± 1.5 | 5.1 ± 3.7 | 4.1 ± 3.5 |

**Table S2.** Cell motility measured by time lapse video of *D. discoideum* expressing different domains of CSP and (I27)4 in presence of 0 µM and 400 µM cAMP and the corresponding cell stiffness measured by AFM indentation assays. The data shown are average ± SD.
